# Neuronal lipofuscinosis caused by Kufs disease/CLN4 *DNAJC5* mutations but not by a CSPα/DNAJC5 deficiency

**DOI:** 10.1101/2023.05.10.540177

**Authors:** Santiago López-Begines, Ángela Lavado-Roldán, Cristina Mesa-Cruz, Fabiola Mavillard, Nozha Borjini, Vera I. Wiersma, Carolina Aguado, Rafael Luján, Wiep Scheper, José L. Nieto-González, Rafael Fernández-Chacón

## Abstract

Kufs disease/CLN4 is an autosomal dominant neurodegenerative disorder that affects young adults, caused by mutations in the DNAJC5 gene that encodes the synaptic vesicle co-chaperone Cysteine String Protein α (CSPα/DNAJC5). The Leu115Arg and Leu116Δ mutations in humans are known to independently cause the disease, although the underlying mechanisms are unknown. To investigate the disease mechanisms in vivo, we generated three independent mouse lines overexpressing different versions of CSPα/DNAJC5 under the neuron-specific Thy1 promoter: wild-type (WT), Leu115Arg, and Leu116Δ. Mice expressing mutant CSPα/DNAJC5 are viable and do not show any significant increase in morbidity or mortality. However, we observed the presence of pathological lipofuscinosis in the mutants, indicated by autofluorescent punctate structures labeled with antibodies against ATP synthase subunit C, which were absent in the WT transgenic line. Additionally, transmission electron microscopy revealed intracellular structures resembling granular osmiophilic deposits (GRODs), observed in Kufs disease patients, in the mutants but not in non-transgenic controls or the WT transgenic mice. Notably, conventional, or conditional knockout mice lacking CSPα/DNAJC5 did not exhibit any signs of increased lipofuscinosis or GRODs. Our novel mouse models thus provide a valuable tool to investigate the molecular mechanisms underlying Kufs disease/CLN4. We conclude that DNAJC5 mutations cause neuronal lipofuscinosis through a cell-autonomous gain of a novel but pathological function of CSPα/DNAJC5.

## INTRODUCTION

Neuronal ceroid lipofuscinosis (NCLs) are inherited neurodegenerative diseases that normally appear in children and young individuals. CLN patients generally suffer from severe neurological symptoms such as seizures, dementia, blindness, and motor disorders that lead to premature death caused by mutations in 13 different genes (termed *CLN1-14*)[7]. Typically, those mutations are loss of function mutations in genes that encode a variety of proteins involved in lysosomal function, such as proteins involved in the trafficking of lysosomes, lysosomal integral membrane proteins and lysosomal enzymes, among others [9, 35, 47]. Those mutations may impact not only neuronal but also microglial function[50]. Autosomal-dominant adult-onset neuronal ceroid lipofuscinosis (CLN4), originally known as Kufs disease, is a severe form of NCL with a symptomatic onset between 25 and 46 years and characterized by generalized seizures, movement disorders, cognitive deterioration, and progressive dementia [5, 18, 26, 28, 33]. The neuronal accumulation of autofluorescent material and the ultrastructural detection of granular osmiophilic deposits (GRODs) are the major neuropathological hallmarks of the disease[5, 18]. Kufs disease/CLN4 is caused by mutations in *DNAJC5* that encodes the synaptic co-chaperone Cysteine-String Proteinα (CSPα/DNAJC5) [2, 8, 25, 38, 52]

CSPα/DNAJC5 is a synaptic vesicle protein that belongs to the heat shock protein-40 (Hsp40) family of molecular co-chaperones. CSPα/DNAJC5 operates through a chaperone complex that includes heat-shock-cognate-70 (Hsc70) and small glutamine-rich tetratricopeptide containing protein-A (SGTA)[51]. Such a trimeric complex acts as a chaperone for the SNARE complex and it is essential to maintain the stability of SNAP25 at synapses [49]. Neurodegeneration and early lethality occur in Drosophila and mice without CSPα/DNAJC5 [17, 60]. Synaptic degeneration induced by absence of CSPα/DNAJC5 is associated with the deficit of SNAP25 but the molecular mechanisms are not well understood yet [49]. Beyond its causative role in Kufs disease/CLN4, CSPα/DNAJC5 has been associated with mechanisms related to other neurodegenerative diseases such as (1) Parkinson’s disease, through the cooperation with α-synuclein as a SNARE complex chaperone [11] and (2) as a mediator of prion-like propagation of key proteins involved in neurodegeneration such as tau, α-synuclein and TDP-43 [13, 19, 29, 55, 56]. In any case, the molecular mechanisms by which the mutant versions of CSPα/DNAJC5 cause Kufs disease/CLN4 are not well understood yet. It has been demonstrated that the two mutations causing Kufs disease/CLN4, Leu115Arg and Leu116Δ, localized at the cysteine string domain, interfere with normal palmitoylation [4, 21, 38] and lead to the formation of molecular accumulations. Several scenarios have been considered to explain the pathological mechanisms associated with these accumulations, which include loss and/or gain of function mechanisms. The most generally accepted scenario, a dominant-negative loss of function mechanism, considers that the accumulations act by sequestering, and suppressing the function of the WT form of CSPα/DNAJC5 [2, 21, 23, 34]. This view is supported by the reduction of CSPα/DNAJC5 levels detected by immunohistochemistry in patients[38] and the co-oligomerization between WT and mutant forms of CSPα/DNAJC5[21, 34]. That notion has also been supported [33] by the fact that CSPα/DNAJC5 heterozygous knock-out mice are essentially normal, whereas CSPα/DNAJC5 knockout mice die early several weeks after birth [17, 20]. Accordingly, a lower gene dose in heterozygous patients would not be pathological in contrast to a severe loss of CSPα/DNAJC5 via a dominant-negative mechanism [33]. The gain of function scenarios consider different possibilities: (1) the mutant accumulations might cause cellular toxicity by recruiting and sequestering other, so far unknown, key cellular proteins [21], (2) the mutant forms themselves maintain their functional co-chaperone properties but they undergo a gain of function evidenced by an increased tendency to form oligomers[59] and (3) the pathological phenotypes are due to a hypermorphic mutation that has a toxic effect mediated by an increasing, so far unknown, activity of CSPα/DNAJC5 mutant forms[24]. Most of the results supporting gain of function phenotypes are based on studies derived from human samples or cell culture studies, except a recent study in a Drosophila model in which the genetic reduction of endogenous levels of CSPα/DNAJC5 ameliorated the pathological phenotype induced by CSPα/DNAJC5 mutant forms[24]. Accordingly, the neurotoxicity of the mutants may be at least in part due to an impairment of lysosomal function by CSPα/DNAJC5 accumulations [24, 59] that accumulate on prelysosomal endosomes [24]. Interestingly, another pathological mechanism proposed for Kufs disease/CLN4 is an aberrant change in the palmitoylation of lysosomal and synaptic proteins caused by a decreased enzymatic activity of the depalmitoylating enzyme palmitoyl-protein thioesterase (PPT1/CLN1)[23]. In any case, although the investigation of pathological mechanisms and development of therapeutical approaches for others CLNs have enormously benefited from mammalian animal models, genetic models in mammals to investigate Kufs disease/CLN4 mechanisms are currently not available.

Here, we have generated independent transgenic mouse lines that express in neurons different versions of CSPα/DNAJC5: WT, Leu115Arg or Leu116Δ. Interestingly, the transgenic mice expressing the mutant forms, clearly develop the typical autofluorescent punctate staining characteristic of lipofuscinosis and GROD-like structures previously described in patients. In addition, lipofuscin engulfing by microglia is also detected in mutant mice. In contrast, lipofuscinosis does not appear in CSPα-WT transgenic mice. In addition, lipofuscinosis is absent in conventional CSPα/DNAJC5 KO mice and, furthermore, it is not detected in aged conditional CSPα/DNAJC5 KO lacking the protein specifically in glutamatergic neurons. Our results indicate that the lack of CSPα/DNAJC5 does not cause lipofuscinosis in mice, whereas the presence of pathological mutant forms in neurons causes lipofuscinosis by a cell-autonomous mechanism. We conclude that the main pathological mechanism driving Kufs disease/CLN4 is likely mediated by aberrant gain-of-function interactions of CSPα/DNAJC5 mutants with so far unknown proteins and not by a dominant-negative mechanism suppressing the function of endogenous non-mutated CSPα/DNAJC5.

## RESULTS

### Expression of CLN4 CSPα mutant forms in transgenic mouse brains

Since CLN4 is an autosomal-dominant disease, we considered to drive the expression of CLN4 CSPα/DNAJC5 mutant forms in neurons over a WT background as a potentially useful strategy to generate CLN4 animal models. We used the Thy-1 promoter to induce the transgenic expression in neurons (Fig. 1 A). The expression of the GFP-CSPα-WT, GFP-CSPα-L115R and GFP-CSPα-L116Δ transgenes was detected in multiple regions of the brain and we selected three transgenic mouse lines with comparable expression patterns in hippocampus and cortex (Fig. 1 B). We studied the transgenic expression patterns and levels in 15 months old (Fig. 1) and in 8 months old mice (Supplementary Fig. 1) and found similar results at both ages. The expression of the GFP-CSPα-WT transgene turned out to be better detectable than the expression of the CLN4-mutant forms, especially when compared with the expression of CSPα-L115R (Fig. 1 B). This was, for example, evident at the hippocampal mossy fibers (Fig. 1 B). To further investigate protein expression levels, we analyzed by immunoblot the levels of endogenous CSPα/DNAJC5 and transgenic proteins (Fig. 1 C). The levels of endogenous CSPα/DNAJC5 were similar in the three transgenic lines and in the non-transgenic controls (Fig. 1 C). We did not observe differences in the levels of the C-subunit of mitochondrial ATPase (ATP5G), that is characteristic of lipofuscin bodies [27, 39, 48]. The expression levels of the transgenic proteins, however, varied among the three lines, with GFP-CSP-L115R having the lowest, and GFP-CSP-L116Δ the highest expression. Remarkably, in CLN4 mutants, especially in the L115R mutant, we found the presence of high molecular weight oligomers of CSPα/DNAJC5, likely corresponding to the CSPα/DNAJC5 accumulations previously described [4, 21, 24, 34, 59](Fig. 1 C, Supplementary Fig. 1).

**Figure 1.**
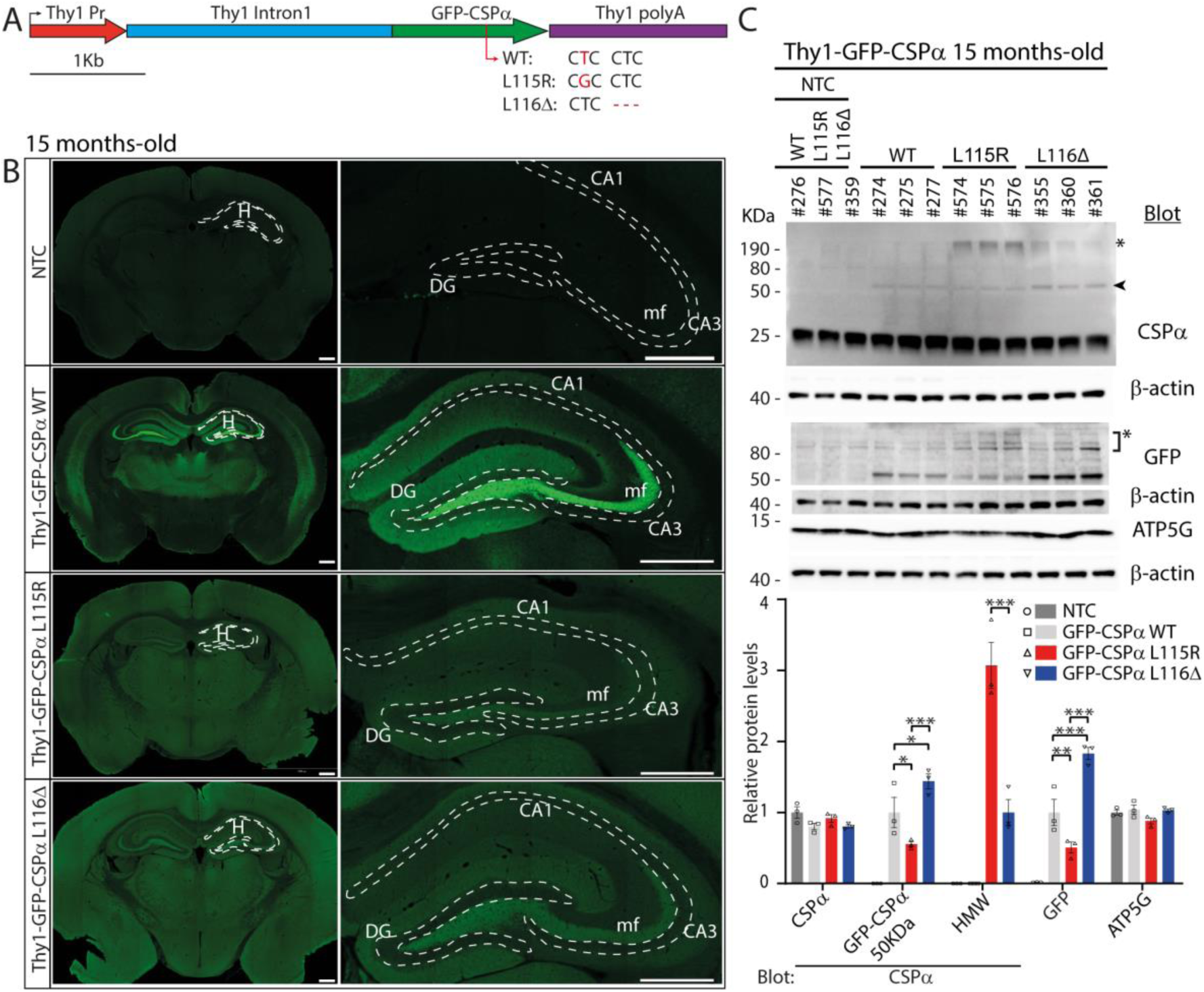
Transgenic expression of wild-type and CLN4 mutant forms of CSPα/DNAJC5 in mouse brain. **A**. Genetic strategy to drive the neuronal expression of GFP-CSPα-WT and CLN4 mutants GFP-CSPα-L115R and GFP-CSPα-L116Δ under the neuron-specific promoter Thy1. **B**. Representative epifluorescence images of GFP immuno-staining demonstrate widely distributed expression of all transgenes in brain in 15 months old mice. General stronger transgene expression of GFP-CSPα WT especially evident at hippocampal mossy fibers. No signal is detected in non-transgenic control (NTC) mice. Scale bar 500µm. **C**. Transgenic proteins detected by western blot of hippocampal extracts from 15 months old mice. Numbers indicate mouse ID number. Upper blot, endogenous CSPα/DNAJC5 is detected in all samples while a band corresponding to GFP-tagged CSPα/DNAJC5 (arrowhead, 50KDa) appears in transgenic samples but not in NTC. High molecular weight species (asterisk *) are detected in mutant transgenic samples, especially in L115R mutant. GFP signal is only detected in transgenic samples. Non-specific band due to GFP antibody (asterisk *). **D**. Levels of selected hippocampal proteins. Relative protein levels normalized to non-transgenic mouse lines, except for GFP quantification that was normalized to GFP levels of the GFP-CSPα-WT transgenic line. Two-way ANOVA with Tukey’s post hoc test (*p<0.05, \*\**p*<0.01, \*\*\**p*<0.001*, ****p<0.0001)*. Quantitative data available at Supplementary Table 1.

### Neuronal lipofuscinosis in CLN4 transgenic mice

Lipofuscinosis is the major hallmark in the brain of Kufs disease/CLN4 patients which is classically found as punctate structures of autofluorescent storage material (AFSM) [5, 33, 48]. We focused on hippocampal slices to investigate with fluorescence microscopy whether the transgenic expression of CLN4 mutants induced the appearance of AFSM (Fig. 2). AFSM is normally excited between 320 and 480nm with an emission spectrum between 460 and 630nm [48]. We used the fluorescein isothiocyanate (FITC) and tetramethylrhodamine isothiocyanate (TRITC) filter sets to examine fluorescence from hippocampal slices with confocal microscopy. Using the FITC filter set, we detected the prominent green fluorescence of GFP at mossy fibers in GFP-CSPα-WT and a weaker general signal in GFP-CSPα-L115R and GFP-CSPα-L116Δ transgenics (Fig. 2 A). Interestingly, only in CLN4 mutants, a punctate pattern of autofluorescence was evident being especially strong in the GFP-CSPα-L115R transgenics at the CA3 region (Fig. 2 A). Such a punctate pattern was also evident using the TRITC filter (Fig. 2 A), suggesting that those structures could be AFSM. We next proceeded to a closer examination at the CA3 region labelling with anti-GFP antibodies (Fig. 2 B, C). In the CLN4 mutants, in which the GFP signal was nevertheless very low (Fig. 2 B), the punctate ATP5G staining was clearly detected at the soma of CA3 pyramidal neurons as perinuclear structures (Fig. 2 C) but not at the mossy fiber terminals stained with antibodies against the synaptic vesicle protein synaptoporin (Fig. 2 C). Some of those punctate structures co-localized with the vestigial GFP staining in the GFP-CSPα-L115R transgenic mice (Fig. 2 B arrowhead). To further investigate if those structures corresponded indeed with AFSM, we labelled the slices with antibodies against ATP5G and found again a clear signal in the CA3 region, especially in the GFP-CSPα-L115R transgenic (Fig. 2 D). The ATP5G signal was absent at the mossy fibers in all the mouse lines, including the GFP-CSPα-WT line (Fig. 2 D). The localization of ATP5G (Fig. 2 D) was highly correlated with the autofluorescence signal detected through the 561-laser line channel (R-Pearson’s correlation coefficient 0.61; see Supplementary Table 1). This was consistent with the notion that ATP5G, likely with other mitochondrial proteins, yields the autofluorescence originated from AFSM.

**Figure 2.**
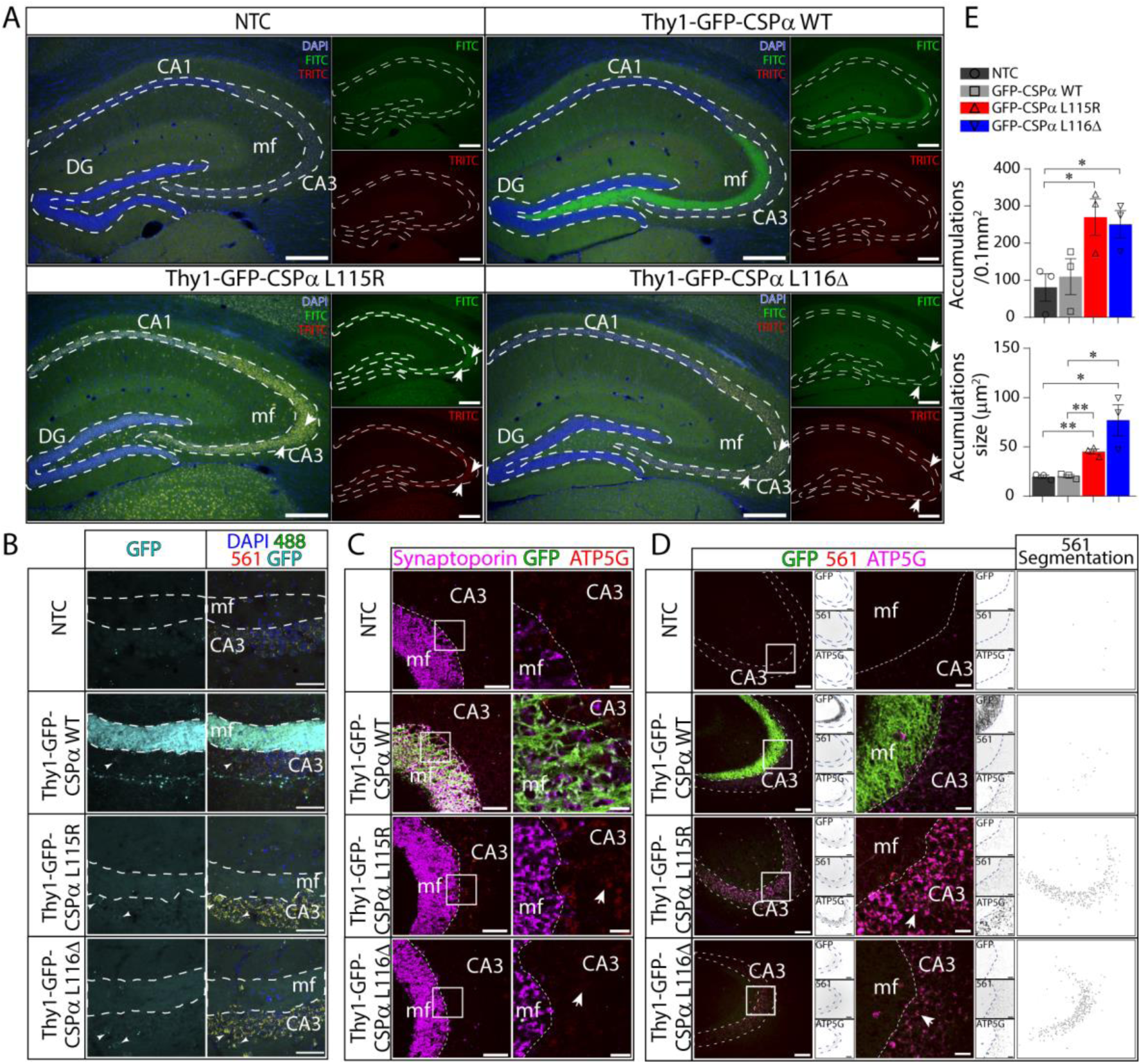
Neuronal lipofuscinosis in Thy1-GFP-CSPα-L115R and Thy1-GFP-CSPα-L116Δ transgenic mice. **A.** Representative merged epifluorescence images of mouse hippocampal slices from non-transgenic control (NTC) and GFP-CSPα-WT, GFP-CSPα-L115R and GFP-CSPα-L116Δ transgenic mice at 15 months-old stained with DAPI (blue). Using the FITC filter set, the overlapping fluorescence signals (green) coming from GFP fluorescence and autofluorescence are collected. By using the TRITC filter set, only autofluorescence signal is collected (red and yellow). Autofluorescence puncta are evident in transgenic mice expressing CLN4 CSPα/DNAJC5 mutations, especially at the pyramidal cell layers (arrows). Scale bar 200µm. **B.** Merged confocal images of mossy fibers at the CA3 region in hippocampal slices stained with antibodies against GFP (cyan) and DAPI (blue). Autofluorescence signal (488 nm and 561 nm laser channels) (yellow) is evident in pyramidal neurons at the CA3 region in CLN4 mutants. Scale bar 50µm. **C.** Confocal images from hippocampal sections stained with anti-ATP5G (red), anti-GFP (green) and anti-synaptoporin (magenta) antibodies. GFP only detected in GFP-CSPα-WT mice at mossy fibers. ATP5G clearly detected at CA3 postsynaptic cells in GFP-CSPα-L115R mice. Scale bar left 50µm.Right 10µm. **D.** As in C plus the collection of autofluorescence through 561 nm laser channel. Autofluorescence and ATP5G signal detected in CLN4 mutants. Small square panels located at the right side of every individual panel respectively display, from top to bottom: the GFP, the 561 nm laser channel and the ATP5G signals. White panels at the right show images resulting after mask segmentation of 561 nm laser channel used for quantification of autofluorescent spots (see Material and Methods). **E.** Significant differences are observed in accumulations size and density. Mean±SEM. N=3 animal/genotype. Unpaired t-test *p<0.05, **p<0.01. Scale bars: left 100µm, right 25µm. Scale bar 50µm. Quantitative data available in Supplementary Table 1.

Strikingly, the prominent GFP signal in mossy fibers in the GFP-CSPα-WT transgenic did not translate into obvious autofluorescence in the mossy fibers terminals or in the soma of CA3 pyramidal neurons. In addition, we could not detect any significant autofluorescence in the non-transgenic mice used as negative controls. The signs of lipofuscinosis were more evident in the GFP-CSPα-L115R than in the GFP-CSPα-L116Δ transgenic mice and in mice older than 4 months, with the highest accumulation in aged 15 months old mice (Supplementary Fig. 2). In addition, to avoid any confounding effect of fluorescence signals we demonstrated the existence of ATP5G accumulations by immunohistochemistry and bright field microscopy (Supplementary Fig. 3).

Next, to investigate the origin of autofluorescence we searched for the presence of granulovacuolar degeneration bodies (GVBs) [54]. Sections were immunolabeled with antibodies against casein kinase 1 δ (CK1δ) and phosphorylated protein kinase R (PKR)-like endoplasmic reticulum kinase (pPERK) that, as previously reported, are useful markers to detect GVBs in the human brain, as well as in mouse and cell [54]. Neither CK1δ (Supplementary Fig. 4) nor pPERK immunolabeling showed GVBs in any of the genotypes. Furthermore, immunolabeling with an antibody against the lysosomal and GVB membrane marker lysosomal integral membrane protein-2 LIMP2 did not reveal clear differences between genotypes (Supplementary Fig. 4). Therefore, the punctate autofluorescence phenotype of mutant GFP-CSPα mice was not reflected by an obvious lysosomal phenotype nor related to the presence of GVBs. We concluded that CSPα/DNAJC5 CLN4 mutations do not induce hippocampal GVB formation.

Altogether, these observations suggest that the autofluorescence is specifically detected only in CLN4 mutant transgenic mice, even under a rather low level of transgenic protein expression compared to the levels of endogenous CSPα/DNAJC5. Autofluorescence likely corresponds to age-dependent pathological lipofuscinosis at the neuronal soma with accumulation of a mitochondrial marker, rather than lysosomal proteins. To further investigate the origin of the autofluorescence we next used electron microscopy to explore ultrastructural intracellular details.

### Granular osmiophilic deposits-like structures in neurons of CLN4 transgenic mice

We examined the soma of hippocampal pyramidal neurons at the CA3 region in 8 months old mice of different phenotypes (Fig. 3 A, left). In non-transgenic control mice, we found cytosolic round compound structures generally composed by a translucent sphere, likely a lipid droplet, attached to darker membrane-bound structures filled with a moderated electron dense material including linear profiles and some very dark dots (Fig. 3 A, left). The same type of structures was found in samples from the GFP-CSPα-WT mice. In contrast, we did not find those structures in GFP-CSPα-L115R mice. Instead, the structures that we found in these mutant mice were much more electron dense, apparently formed by accumulated material and with less lipid-droplets associated to them. High electron density structures may indicate a high concentration of macromolecules or higher binding of heavy metals such as osmium. Indeed, the structures strongly resembled the GRODs found in Kufs disease/CLN4 patients [1, 5, 18, 38, 52], so we named them GROD-like structures. One possibility is that the structures found in the mutants were not pathological inclusions, but they reflected, instead, the advanced stage of an otherwise physiological age-dependent accumulation of lipofuscin that should be found in aged control mice. To further investigate this possibility, we analyzed aged 15 months-old mice (Fig 3 A, right). The results were similar to those found in younger 8 months-old mice. We could not find GROD-like structures in non-transgenic control and GFP-CSPα-WT transgenic aged mice in contrast to the evident presence in GFP-CSPα-L115R transgenic mice at both ages (Fig 3 A, right). This observation suggests that GROD-like structures do not appear because of physiological aging in control mice, and they are instead a specific feature of the CLN4 transgenic mutant mice. Although the GROD-like structures were most evident in GFP-CSPα-L115R mice at both ages studied, they were also detected in GFP-CSPα-L116Δ mice but to a lesser extent (Fig. 3 A). Indeed, the number of lipofuscin particles, including GROD-like structures, were significantly higher in GFP-CSPα-L115R transgenic mice compared to controls, but not in GFP-CSPα-L116Δ transgenic mice (Fig. 3 B).

**Figure 3.**
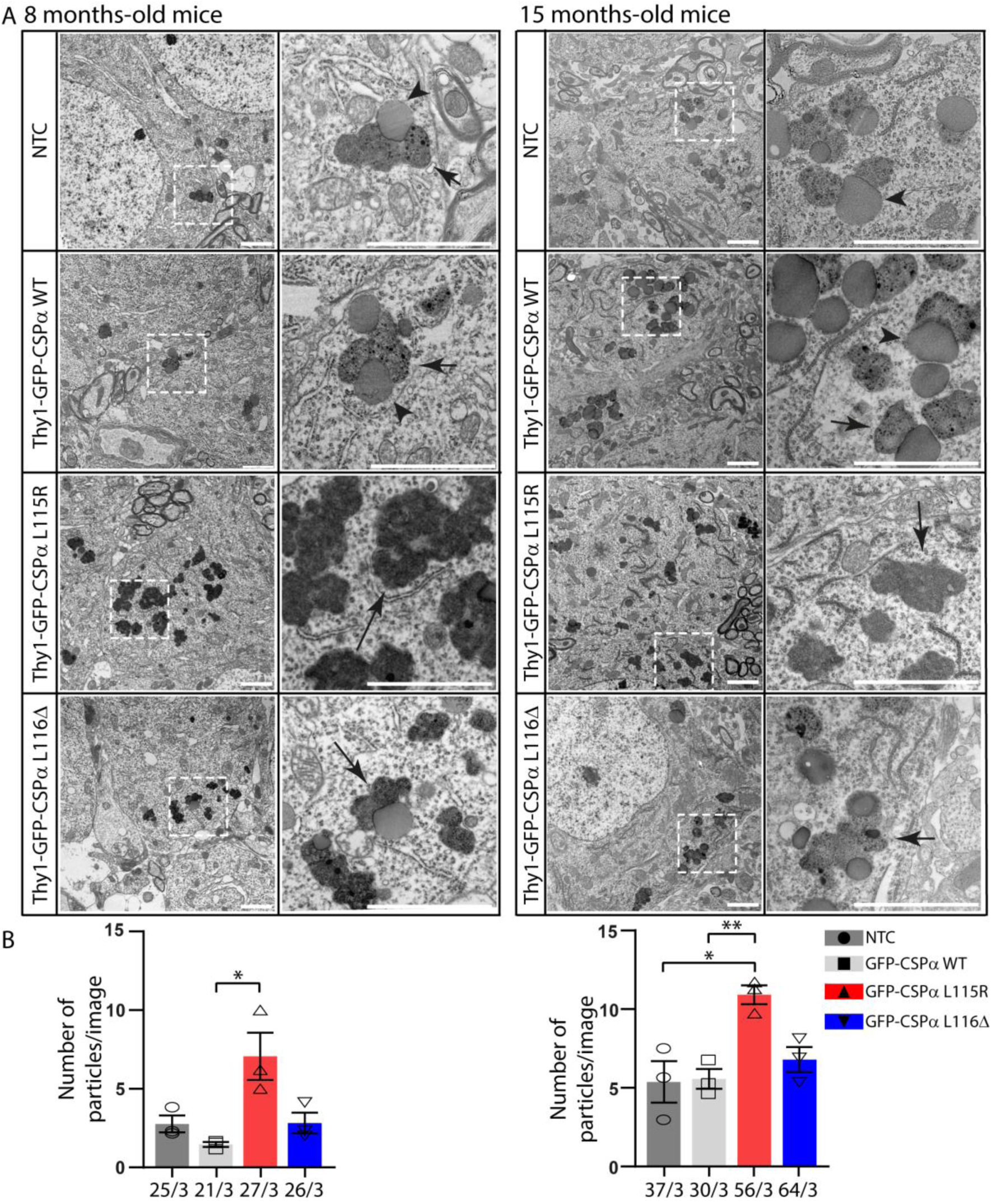
Granular osmiophilic deposits-like (GROD-like) structures in the somata of CA3 pyramidal neurons of CLN4 transgenic mice at 8 and 15 months age. **A**. Transmission electron microscopy analysis reveals normal lipofuscin characterized by lipid droplets associated to rather clear structures with dark puncta (short arrows) in control mice (NTC and Thy1-GFP-CSPα-WT) in contrast to dark GROD-like structures (long arrows) found in mutant mice (Thy1-GFP-CSPα-L115R and Thy1-GFP-CSPα-L116Δ). Pictures at right columns are magnification of selected areas at the left columns. **B**. Lipofuscin and GROD-like structures were all quantified and plotted as particles per image revealing an increased number in the GFP-CSPα-L115R mutant at both ages studied (8 months on the left and 15 months on the right). 3 animals were studied per genotype, unpaired T-test *p<0.05, **p<0.01. Scale bar 2µm. NTC (non-transgenic control).

### Microglia engulfing of lipofuscin in CLN4 mutant mice

Since microglia alterations have been previously reported in tissue of CLN4 patients[3] and in mouse models of NCLs [45, 46, 57, 58], we investigated the status of microglia in CLN4 mutant transgenic mice. Microglia are continuously extending and retracting their processes to survey the brain [12, 37] and target damaged cellular structures to envelop them, a process normally associated with changes in microglia morphology [31, 40]. We used antibodies against the ionized calcium-binding adapter molecule 1 (Iba1) to study microglia with confocal microscopy at the CA3 region of GFP-CSPα-WT, GFP-CSPα-L115R, GFP-CSPα-L116Δ transgenic and non-transgenic mice (Fig. 4). Microglia were equally abundant in CLN4 mutant and control but surrounded by autofluorescence storage material in CLN4 mutant mice (Fig. 4 A). We used Imaris 3D reconstruction to quantify the morphological properties of microglia, focusing on (1) the ramification of their processes, the size of cell bodies (Fig. 4 A) and to (2) the spatial relationship between microglia and lipofuscin visualized as autofluorescence storage material (Fig. 4 A, B). First, we measured the area of the cell body, the total area occupied by microglia, and the processes length per cell in GFP-CSPα-L115R (Supplementary Fig. 5) and GFP-CSPα-L116Δ mice (Supplementary Fig. 6) and compared them with the values obtained from GFP-CSPα-WT and non-transgenic controls (both 15 months of age) at different ages (1, 4, 8 and 15 months of age). We found only a moderate increase in cell body size in the microglia of GFP-CSPα-L115R mice and a decrease in the length of the microglial processes of GFP-CSPα-L115R and GFP-CSPα-L116Δ mice (Supplementary Fig. 5 and 6).

**Figure 4.**
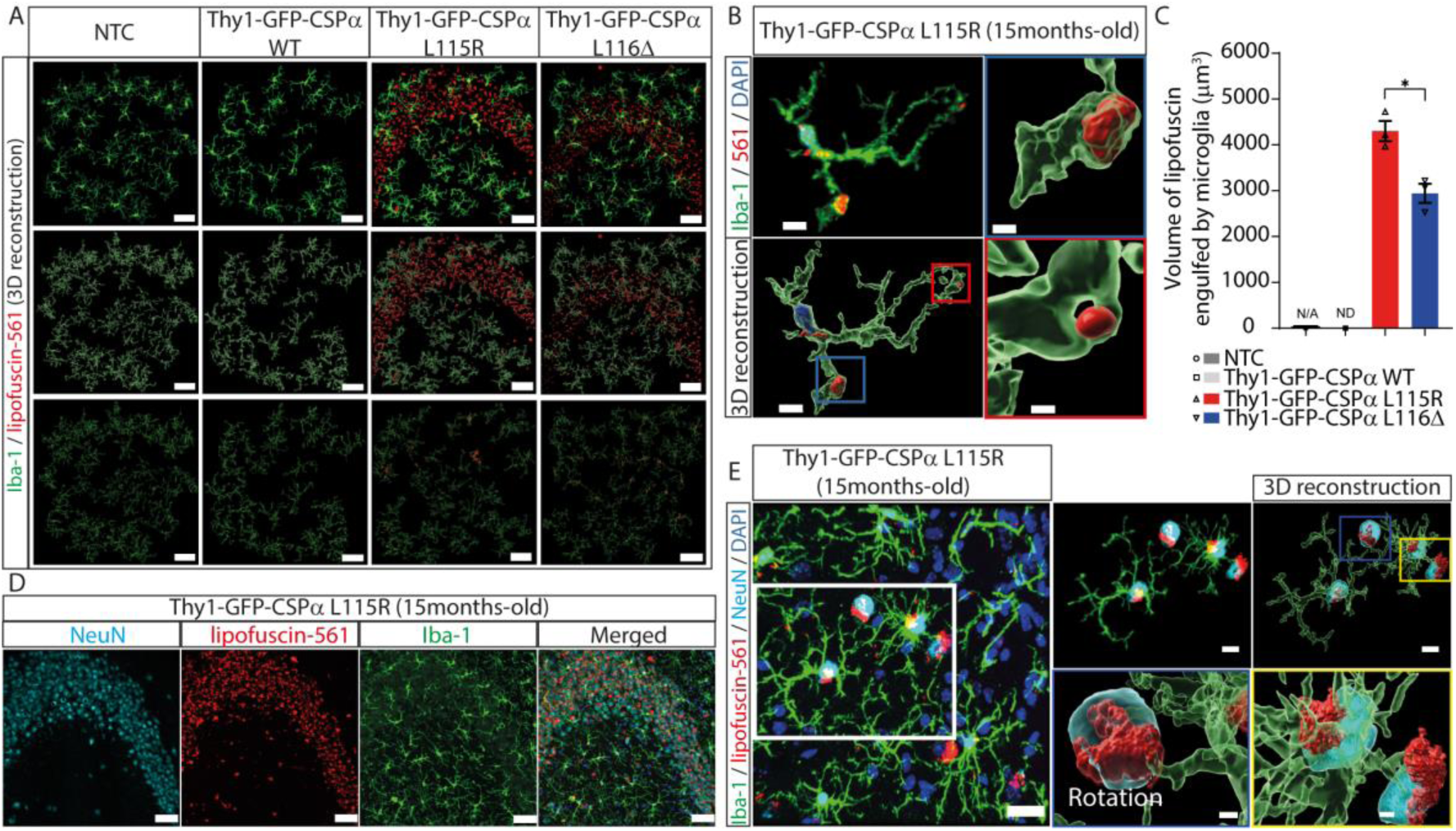
Microglia engulf lipofuscin and neurons in Thy1-GFP-CSPα-L115R and Thy1-GFP-CSPα-L116Δ transgenic mice. **A.** First row, representative images of immunostaining of microglia (Iba-1, green) and lipofuscin (red, autofluorescence at 561nm using the TRITC filter) at the CA3 region of Thy1-GFP CSPα L115R of 15 months old mutant mice. **Second row**, Imaris 3D reconstruction. **Third row**, Imaris 3D reconstruction of lipofuscin engulfed by microglia. Scale bar 50µm. **B.** Imaris 3D reconstruction of one microglia cell of Thy1-GFP CSPα L115R mutant mice at 15 months old showing engulfed lipofuscin particles by microglia. Scale bars 8µm, 2µm (red and blue boxes) **C.** Quantification of volume of lipofuscin engulfed by microglia in control (NTC) and GFP-CSPα-WT, GFP-CSPα-L115R and GFP-CSPα-L116Δ transgenic mice. Data were presented as mean ± SEM, Two-way ANOVA with Tukey’s post hoc test (**p<0.01, ***p<0.001 ****p<0.0001), N=3 mice/group, 3 images/mouse. At least 25 microglia cells/per mouse were analyzed to get the processes length of microglia data. **D.** Immunostaining of microglia (Iba-1, green), lipofuscin (red, autofluorescence at 561nm), neurons (NeuN, cyan) and nuclei (DAPI, blue) of 15 months old Thy1-GFP CSPα L115R mice. Scale bar 50µm. **E.** Representative images of Imaris 3D reconstruction showing the close interaction between microglia and neurons with lipofuscin (blue square) and neurons with lipofuscin that is engulfed by microglia (yellow square) in Thy1-GFP CSPα L115R mice. Scale bars 20µm,10µm (white box), 1µm (blue and yellow boxes). Part of the data set used to generate panels A and C in Figure 4 (15month-old mice data) were also used for Supp. Fig. 7 and 8. NTC and Thy1-GFP-CSPα WT data set (both from 15 month-old mice) are the same in Fig. 4 and in Suppl. Fig. 7.

These morphological changes are somehow like those occurring in the interleukin 1β-releasing microglia [31, 40]. In addition, we observed that, although the majority of autofluorescence storage material was located around microglia (Fig. 4 A and Supplementary Fig. 5 and 6), there were detectable autofluorescence particles within or engulfed by microglial processes in CLN4 mutants (Fig. 4 B), being more prominent in GFP-CSPα-L115R than in GFP-CSPα-L116Δ mice at 8 and 15 months of age (Supplementary Fig. 7 and 8).

Since lipofuscin was not detectable in control mice we could not see microglia engulfing events either. We could not distinguish if the autofluorescent material engulfed by microglia corresponded to lipofuscin released to the extracellular space or corresponded, instead, to lipofuscin contained within a neuron. In any case, upon labeling neurons with anti-NeuN, we observed microglia embracing neuronal bodies containing lipofuscin, so we concluded that microglia could at least engulf neurons containing lipofuscin, as previously described[6, 32].

Altogether, our data reveal that the transgenic expression of mutant forms of CSPα/DNAJC5 in neurons is enough to cause neuronal lipofuscinosis in mice that is similar to the pathological characteristics of Kufs disease/CLN4 in humans[5, 18]. Importantly, these findings identify our mice as novel and unique *in vivo* models to investigate the molecular basis of lipofuscinosis in Kufs disease/CLN4. A key question arising is which is the mechanism underlying the genesis of lipofuscinosis. CSPα/DNAJC5 mutated forms might induce the pathology by (1) a loss of function mechanism as a secondary decrease in the levels of endogenous non-mutated CSPα/DNAJC5, (2) a hypermorphic gain of function mechanism as previously shown in Drosophila[24], and (3) a mechanism by which the CSPα/DNAJC5 mutated forms would sequester specific proteins required for an as of yet unknown biochemical pathway that prevents pathological lipofuscinosis. These scenarios do not necessarily exclude one another, and we proceeded to investigate which one of them could be the culprit for pathological lipofuscinosis.

### Acute and long-term genetic removal of CSPα/DNAJC5 does not cause lipofuscinosis in mice

If lipofuscinosis were induced upon a decrease of CSPα/DNAJC5 levels, then its presence should be evident in the neurons of CSPα/DNAJC5 KO mice. These mice suffer from early lethality and barely reach one month of postnatal age, so the oldest mice we could use for this study were 30 days old mice to be compared with CSPα/DNAJC5 WT control littermates. As with the transgenic mice, we used electron microscopy to analyze the somata of pyramidal neurons at the hippocampal CA3 region (Fig. 5). Neither in CSPα/DNAJC5 KO nor in littermate control mice we detected any GROD-like structure. We only detected membrane-bound structures filled with moderate electron dense content including linear profiles and dark spots (Fig. 5 A, C). They were like those found in 8 months-old and 15 months-old non-transgenic controls and GFP-CSPα-WT transgenic mice but without lipid-droplets attached. Those observations were solid enough to conclude that the mere removal of CSPα/DNAJC5 does not translate into the immediate generation of lipofuscinosis. Nevertheless, those observations did not rule out that a partial, but long term, reduction of CSPα/DNAJC5 levels in 8 and 15 months-old transgenic mutant mice could be the cause of lipofuscinosis. To further explore this possibility, we examined 8 months-old heterozygous CSPα/DNAJC5 KO and WT littermates to investigate if the gene dosage reduction of CSPα/DNAJC5 constitutively found in heterozygous mice (Supplementary Fig. 9) was enough to build up lipofuscinosis. The electron microscopy analysis, however, only detected the same ultrastructural features previously found in non-transgenic controls and GFP-CSPα-WT transgenic mice without any evidence of GROD-like structures (Fig. 5 B, C). Still these results could be explained if the decrease of ∼35% in CSPα/DNAJC5 levels in the heterozygous mice (Supplementary Fig. 9) was not as strong as in the transgenic mutants. To definitely check if a long-term severe reduction of CSPα/DNAJC5 in adulthood would be enough to induce lipofuscinosis, we searched for signs of lipofuscinosis in CaMKIIα^CreERT2^Ai27D:Dnajc5^flox/-^ conditional KO mice by comparing them to CaMKIIα^CreERT2^:Ai27D:Dnajc5^flox/+^ littermate control mice. The CaMKIIα^CreERT2^:Ai27D:Dnajc5^flox/-^ conditional KO mice fed with tamoxifen (TMX) (Fig. 6 A) were viable and did not show an obvious increase in morbidity and only an increase in mortality at old ages (61% of CaMKIIα^CreERT2^:Ai27D:Dnajc5^flox/-^ survived up to 17 months). A deeper characterization of these mice will be described elsewhere (Mesa-Cruz et al., in preparation). The analysis of protein expression in hippocampal sections using fluorescently labeled antibodies clearly demonstrated the absence of CSPα/DNAJC5 at the mossy fibers and at the Schaffer collaterals in 3 months-old mice one month after the TMX-diet was completed (Supplementary Fig. 10). In addition, the neurons in which cre-recombinase was active were identified as red fluorescent neurons because of the expression of the reporter td-tomato. Therefore, these mice were ideal to investigate if lipofuscinosis appears associated to the long-term absence of CSPα/DNAJC5 during adulthood and during aging. We investigated cohorts of mice at 8, 16 and 22 months after the TMX-diet was removed (Fig. 6). As expected, CSPα/DNAJC5 was absent from hippocampal glutamatergic neurons of CaMKIIα^CreERT2^:Ai27D:Dnajc5^flox/-^ mice except in the mossy fibers terminals in which we found sparse CSPα/DNAJC5 staining (Supplementary Fig. 10) that curiously was not present in younger 3 months-old mice. We interpreted that the presence of CSPα/DNAJC5 came from the mossy-fiber terminals of new granule cells born by adult neurogenesis after TMX discontinuation. As expected, CSPα/DNAJC5 was absent from other neurons that do not undergo adult neurogenesis, such as the CA3 pyramidal neurons, on which our analysis was focused. We stained hippocampal sections with antibodies against the lipofuscinosis marker ATP5G and did not find any difference between CaMKIIα^CreERT2^:Ai27D:Dnajc5^flox/-^ conditional KO mice and controls (Fig. 6 B). Furthermore, we examined the CA3 hippocampal region with transmission electron microscopy and did not find any sign of pathological lipofuscinosis such as GROD-like structures (Fig. 6 C, D). Altogether, our results indicate that the absence of CSPα/DNAJC5 by itself does not cause the pathological lipofuscinosis that, on the other hand, is clearly observed in the transgenic mice expressing the mutant forms of CSPα/DNAJC5 causing Kufs disease/CLN4 in humans.

**Figure 5.**
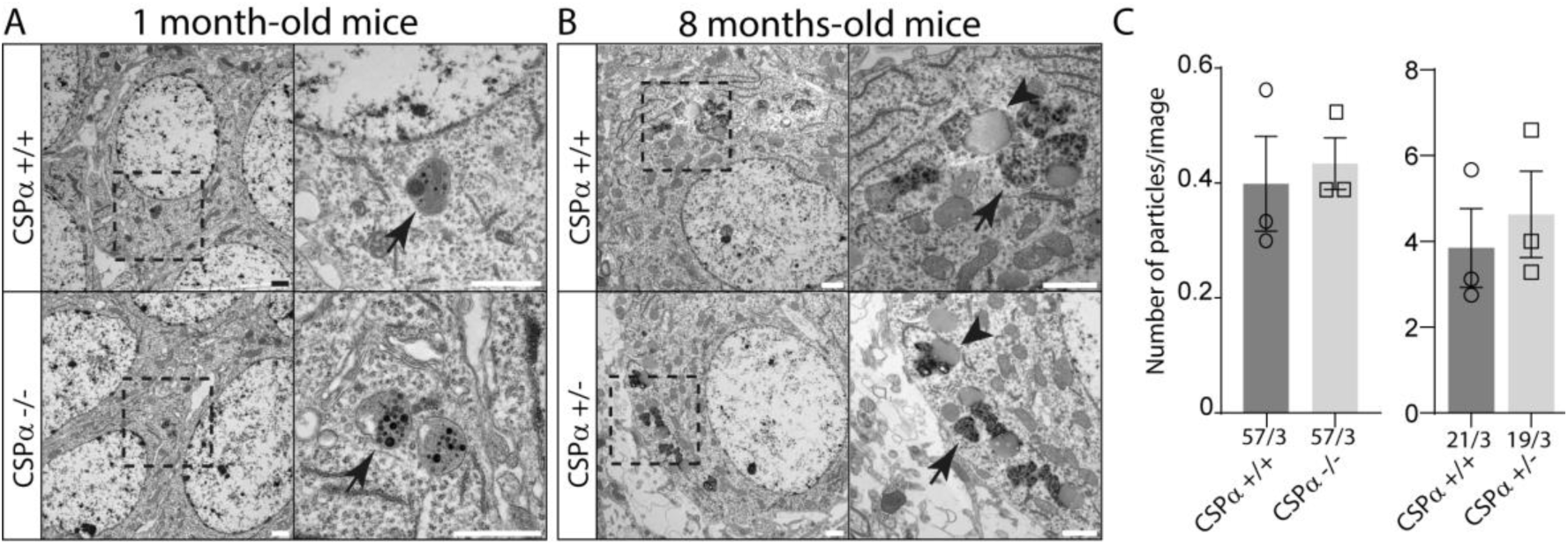
Normal lipofuscin but not GROD-like structures in conventional CSPα/DNAJC5 KO and heterozygous mice. **A**. Transmission electron microscopy analysis reveals that only normal lipofuscin is rarely found at young CSPα/DNAJC5 KO and WT mice at 1-month of age. **B.** Normal lipofuscin, but not GROD-like structures, detected at 8-months-old CSPα/DNAJC5 heterozygous mice. **C.** The number of lipofuscin particles per image in both groups is like control mice, however, the numbers detected in the CSPα/DNAJC5 KO and their controls at 1-month of age. Numbers at the bottom of graph bars indicate number of images/number of mice. Quantitative data available at Supplementary Table 1.

**Figure 6.**
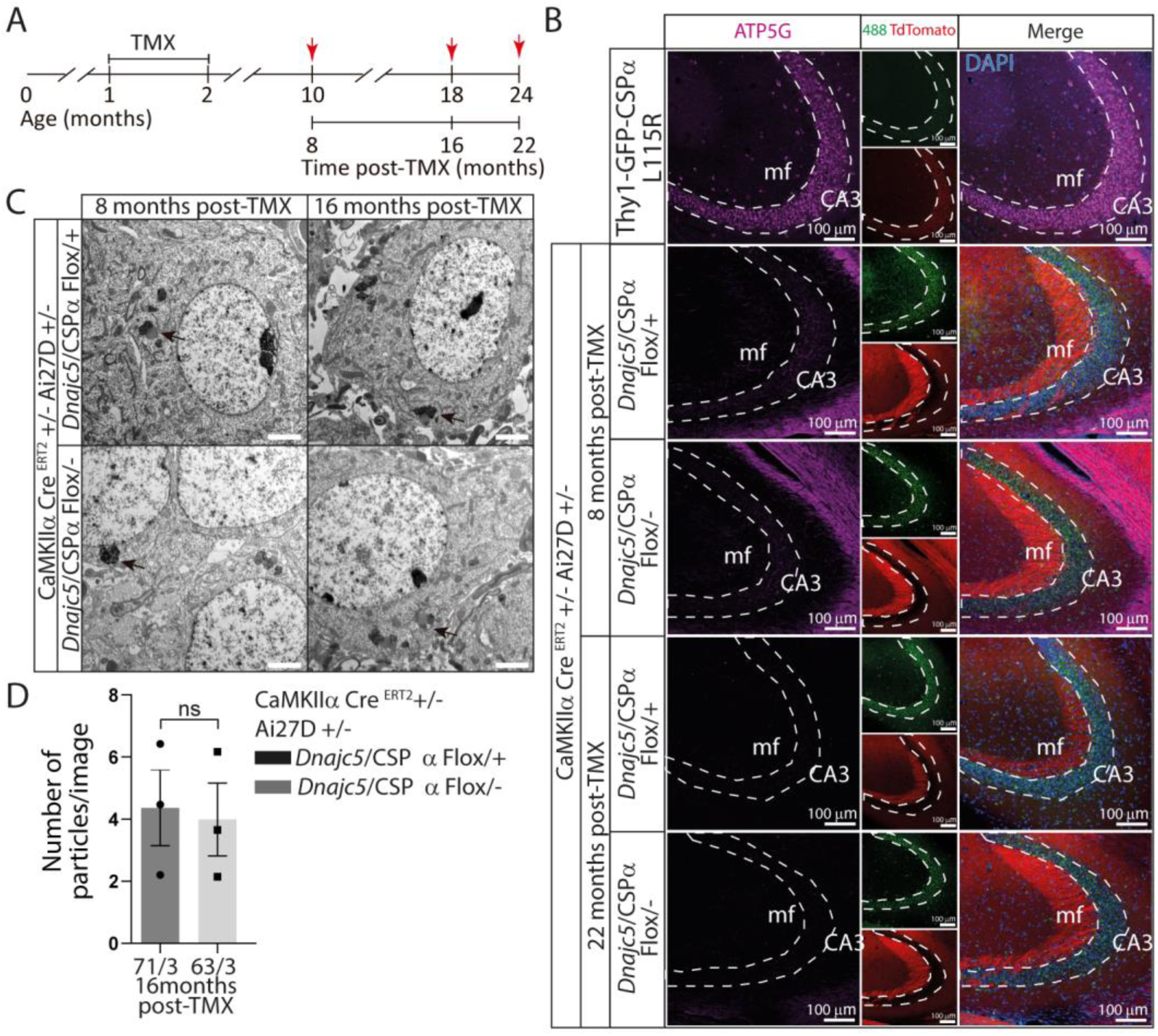
Long term removal of CSPα/DNAJC5 from glutamatergic neurons does not induce pathological lipofuscinosis. **A.** Chronogram of the strategy used for the genetic removal of CSPα/DNAJC5 in control (CaMKIIα^CreERT2^:Ai27D:Dnajc5^flox/+^) and experimental (CaMKIIα^CreERT2^:Ai27D:Dnajc5^flox/-^) mice. Brains were harvested after feeding one or two month old mice with tamoxifen (TMX) (red arrows) for 30 days. Brains of mice fed at two months of age were harvested after 8 months and mice fed at one month of age were harvested after 16 and 22 months post-tamoxifen. **B.** Thy1-GFP-CSPα-L115R mice displays pathological lipofuscinosis at CA3 pyramidal neurons as demonstrated by the immuno-labelling with anti-ATP5G antibodies (magenta) in contrast to mice lacking CSPα/DNAJC5 in hippocampal glutamatergic neurons (CaMKIIα^CreERT2^:Ai27D:Dnajc5^flox/-^) and their controls (CaMKIIα^CreERT2^:Ai27D:Dnajc5^flox/+^) in which anti-ATP5G staining is negative. GFP is detected in green and the reporter Ai27D (channelrhodopsin 2 fused to td-tomato) in red. Scale bar 100µm**. C**. Transmission electron microscopy analysis only detects normal lipofuscinosis in experimental and control mice at 8 and 16 months after tamoxifen treatment. Scale bar 2µm. **D.** The number of lipofuscin particles per image is similar among the experimental and the respective control mice studied at 8-and 16-months post-tamoxifen. Numbers at the bottom of graph bars indicate number of images/number of mice. Quantitative data available at Supplementary Table 1.

### Lipofuscinosis persists upon long-term genetic lowering of CSPα levels in transgenic mice expressing Kufs disease/CLN4 mutations

In Drosophila, the transgenic overexpression of L115R and L116Δ mutations causes lipofuscinosis accompanied by a prominent neurological phenotype [24]. Interestingly, the phenotype is ameliorated when the mutation is expressed on a CSPα/DNAJC5 hypomorphic genetic background in which the genetic dose of endogenous WT CSPα/DNAJC5 is significantly decreased. To test that notion in mammals, we bred heterozygous CSPα/DNAJC5 KO mice, that present a reduction in CSPα/DNAJC5 levels (Supplementary Fig. 9), against the GFP-CSPα-L115R transgenic mice. Upon sequential breeding, we generated GFP-CSPα-L115R transgenic mice that were either homozygous or heterozygous CSPα/DNAJC5 KO mice that were studied and compared to their CSPα-L115R transgenic littermates on a CSPα/DNAJC5 WT genetic background (Fig. 7 A). As previously explained, the early lethality of CSPα/DNAJC5 KO mice allowed studies only in one month old mice that were at the same time GFP-CSPα-L115R transgenic and CSPα/DNAJC5 KO mice. In any case, we could not detect any signs of lipofuscinosis in those mice and the findings were similar to our observations in GFP-CSPα-L115R transgenic mice with a CSPα/DNAJC5 WT background at one month of age (Supplementary Fig. 2). The failure to detect lipofuscinosis in these mice, as we previously observed (Fig. 5 A), is likely due to the youth of the mice because lipofuscinosis only appears in older mutant CSPα/DNAJC5 transgenic mice, usually beyond 4 months of age (Supplementary Fig. 2). Because of that, we next examined 8 months old GFP-CSPα-L115R transgenic CSPα KO heterozygous mice (Fig. 7 B). As expected, we could detect lipofuscinosis in every GFP-CSPα-L115R transgenic mouse examined. Nevertheless, among those mice we could not find any differences between CSPα/DNAJC5 KO heterozygous and CSPα/DNAJC5 WT control mice (Fig. 7 B). These results indicate that, at least in 8 months old mice, a genetic dosage reduction of CSPα/DNAJC5 does not ameliorate pathological lipofuscinosis. Overall, our results indicate that neuronal lipofuscinosis does not occur due to a loss-of-function mechanism, but rather because CLN4 mutations transform CSPα/DNAJC5 in a protein with novel properties that likely interferes with a cellular process through new, but pathological, protein-protein interactions.

**Figure 7.**
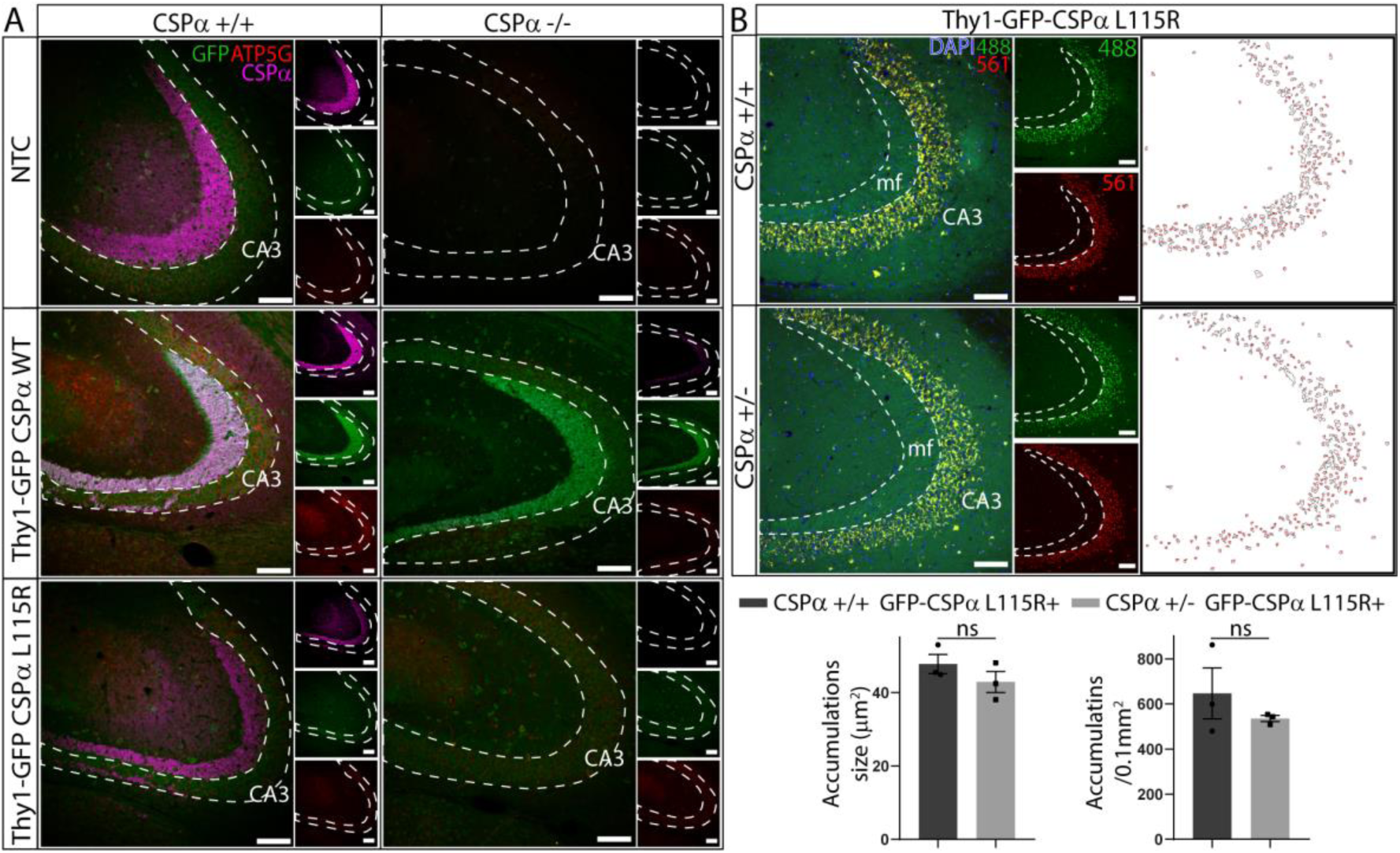
Pathological lipofuscinosis persists upon reducing the endogenous CSPα/DNAJC5 gene dosage. **A.** 1 month old mice expressing the WT (Thy1-GFP-CSPα-WT) and L115R (Thy1-GFP-CSPα-L115R) transgenic versions of CSPα/DNAJC5 in control and CSPα/DNAJC5 KO backgrounds do not show a prominent staining with anti-ATP5G antibodies (red) at the hippocampal CA3 region. CSPα/DNAJC5 (magenta) is detected in controls but it is absent in CSPα/DNAJC5 KO mice. Anti-GFP signal detected in green. Scale bar 100µm**. B.** Pathological lipofuscin is detected as a prominent autofluorescent overlapping signal between the signals coming from the 488nm laser channel (green) and from the 561nm laser channel (red) at the hippocampal CA3 region of Thy1-GFP-CSPα-L115R under CSPα/DNAJC5 WT and heterozygous backgrounds. Anti-GFP signal detected in green and DAPI in blue. Scale bar 100µm. White panels at the right show images resulting after mask segmentation of 561 nm laser channel used for quantification of autofluorescent spots (see Material and Methods). No significant differences (unpaired t-test) are observed in accumulations size and density. Mean±SEM, n=3 animal/genotype. Quantitative data available at Supplementary Table 1.

## DISCUSSION

We have found that transgenic mice expressing the mutations found in patients (L115R and L116Δ) in *DNAJC5* develop neuronal lipofuscinosis with the pathological hallmarks observed in Kufs disease/CLN4 patients and that this lipofuscinosis is not due to the lack of CSPα/DNAJC5. Autosomal dominant Kufs disease/CLN4 is an adult-onset neuronal ceroid lipofuscinosis characterized by a progressing devastating neurodegenerative symptomatology affecting individuals after the third decade of life [26, 28, 33]. The primary cause of the disease, mutations in the DNAJC5 gene, is well stablished, however, the underlying cellular and molecular pathological mechanisms remain elusive. Since the patients bear the mutations in heterozygosis, we used conventional pronuclear injections to drive the expression of the WT and mutant versions of *Dnajc5* fused to GFP. We used the neuronal specific promoter Thy-1 that normally produces a variegated expression in neuronal subpopulations with a moderate expression level ideal to avoid aberrant unphysiological effects caused by excessive overexpression [16]. All the transgenic mice were viable and did not develop any obvious increase in morbidity compared to non-transgenic controls. We focused our attention on the hippocampus and the cerebral cortex as brain areas in which the transgene expression was detected by GFP-fluorescence or by immunofluorescence using antibodies against GFP (Fig. 1). Interestingly, the transgenic WT version of CSPα/DNAJC5 turned out to have a much stronger hippocampal expression, especially in the mossy fibers, compared with the mutant versions, which in general were expressed at lower levels (Fig. 1B). These differences may reflect that the mutant proteins display either low stability and/or undergo conformational changes or accumulation that decrease GFP-fluorescence and/or mask the detection by antibodies. Supporting this notion, it has been reported that palmitoylated monomers of CSPα/DNAJC5 mutants are short-lived compared to WT CSPα/DNAJC5, suggesting that the mutants could have a faster rate of depalmitoylation and/or they are consumed faster in a time-dependent manner into high molecular weight accumulations [14]. Interestingly, the transgenic protein levels detected in hippocampus by western blot were not lower in the mutants compared to the WT transgenic version, yet the levels of all transgenic proteins were much lower than the levels of endogenous CSPα/DNAJC5 (Fig.1C). Strikingly, the mutant proteins induced high molecular weight accumulations that were especially evident in the L115R mutant (Fig.1C). The accumulations contained at least the GFP-fused mutant proteins, however, we do not know if they co-accumulated with the endogenous CSPα/DNAJC5. This biochemical alteration, which was not observed for the WT transgenic protein, is consistent with the previously described increased tendency of the mutant forms of CSPα/DNAJC5 to accumulate [4, 21, 24, 34, 59]. Since a major hallmark of lipofuscinosis is the presence of autofluorescence [1] we searched the hippocampus for autofluorescent storage material and found a remarkable accumulation of punctate autofluorescent structures in the soma of CA3 pyramidal neurons in both mutants, being most pronounced in the L115R mutant (Fig. 2). Two aspects of this observation are remarkable: (1) we did not observe these structures in the non-transgenic control mice and, in addition, the overexpression of the CSPα/DNAJC5-WT transgene did not generate those structures, indicating that the autofluorescence specifically occurred as a result of expression of CSPα/DNAJC5 mutant versions and not by the overexpression of normal CSPα/DNAJC5, (2) the apparently low levels of L115R-CSPα and L116Δ-CSPα proteins detected by immunofluorescence at the CA3 were nevertheless high enough to build-up autofluorescence, reinforcing the notion that the pathogenic effect is rather precisely mediated by the mutations and not because of a non-specific consequence of higher expression levels that, in any case is much lower that the endogenous levels of CSPα/DNAJC5. In addition, we found that the autofluorescent structures were stained with antibodies against ATP5G (also known as mitochondrial ATP synthase subunit C), a protein typically used as lipofuscinosis marker [15, 22]. Remarkably, the electron microscopy analysis revealed the existence of GROD-like structures in the CA3 pyramidal cell soma of mutant mice with the same characteristics of GRODs previously described in brain samples of Kufs disease/CLN4 patients [18, 38]. In non-transgenic control mice and in transgenic mice expressing the CSPα/DNAJC5 WT version, we observed different structures, which were less electron dense, sometimes with internal linear membranous arrays and often associated to a lipid vacuole likely corresponding to physiological age-dependent lipofuscin (Fig. 3). Lipid vacuoles were sometimes found in the L116Δ mutant, but not in the L115R mutant (Fig. 3). This is remarkably consistent with the human pathology of Kufs disease/CLN4 in which the typical clear component of adult lipofuscin is scanty [18, 26]. We could not detect the accumulation or the existence of granulovacuolar degeneration bodies (Supplementary Fig. 4) which are neuron-selective lysosomal structures associated with neurodegeneration and detected by lysosomal markers [53, 54]. The absence hereof suggests that the CSPα/DNAJC5 mutants do not cause the accumulation of lysosomal-related structures, although CSPα/DNAJC5 has been described to be in lysosomes and to be related to lysosomal function and dysfunction [4, 24, 36]. In any case, the mutations might be interfering with lysosomal-mediated autophagy [4, 24] and impairing the degradation of, for example, oxidized mitochondrial proteins that accumulate as autofluorescent stored material. On the other hand, several studies have reported that CSPα/DNAJC5 mediates the extracellular release of neurodegenerative associated proteins [13, 19, 55, 56]suggesting a general role for CSPα/DNAJC5 in misfolding-associated protein secretion (MAPS) [30]. Indeed, it has been proposed that the mutant versions of CSPα/DNAJC5 impair MAPS without interfering with microautophagy [29]. According to that notion, the GROD-like structures we have observed could be caused by a defect in MAPS, although further investigation is required to specifically test that notion in mice. We have detected moderate morphological changes in microglia, consistent with mildly increased microglia activation (Fig. 4), as it has been also reported at the early stage of Kufs disease [3]. Furthermore, we observed microglia engulfing lipofuscin and lipofuscin-containing neurons, which could reflect the physiological phagocytic activity of normal microglia on abnormal neurons as previously described (Fig. 4) [6, 32]. Our lipofuscinosis models in mice complement the existing Drosophila model [24]. The models share key pathological features previously described in post-mortem brain samples of human Kufs disease/CLN4 patients, such as the excessive formation of oligomers and the existence of electron dense structures detectable with electron microscopy [14, 18, 21, 26, 34, 59], which, especially in mice, mirror the autofluorescence and the GRODs found in humans. In flies and mice, the L115R mutation has in general stronger effects than the L116Δ as was also previously observed in human brain samples and neuronal cultures [18, 20]. Interestingly the phenotype in mutant flies is dose-dependent [24]. Flies expressing a single copy of the mutations showed higher levels of oligomers but did not suffer from neurodegeneration or any neurological phenotype which, in contrast, did appear upon the expression of two copies of the mutated gene[24]. We cannot rule out that stronger phenotypes might also occur in mice upon increasing the mutant gene dosage. In any case, our mice open unique opportunities to investigate the temporal cascade of molecular and cellular events that lead to neuronal lipofuscinosis and its relationship with other pathological alterations *in vivo*. An open question in the field is if the Kufs disease/CLN4 phenotype is secondary to a loss of function of CSPα/DNAJC5 due to a dominant negative effect of the mutant forms that would be interfering with endogenous CSPα/DNAJC5[2, 21, 23, 34, 38, 59]. We have investigated this notion studying several CSPα/DNAJC5 knock-out lines. We concluded that the absence of CSPα/DNAJC5 by itself does not cause lipofuscinosis because we do not detect any sign of pathological lipofuscinosis using confocal and electron microscopy in several genetic mouse models such as the CSPα/DNAJC5 knock-out mice at one month of age and CSPα/DNAJC5 heterozygous knock-out mice at eight months of age. However, those models might have limitations. On one hand, the conventional CSPα/DNAJC5 KO mice die very early when they are just one month old, so it could be argued that the mice are still too young to develop lipofuscinosis. On the other hand, although the heterozygous CSPα/DNAJC5 mice were studied during adulthood, it could be argued that the 50% reduction in CSPα/DNAJC5 is not enough to induce lipofuscinosis. However, we overcame those limitations using the CaMKIIα^CreERT2^:Ai27D:Dnajc5^flox^ mouse line in which CA3 pyramidal neurons, among other forebrain glutamatergic neurons, lack CSPα/DNAJC5 for up to 22 months without developing pathological lipofuscinosis. This observation strongly supports the notion that the mere elimination of CSPα/DNAJC5 in neurons is not enough to cause lipofuscinosis. Interestingly, the phenotype observed in Drosophila mutants is a hypermorphic gain of function phenotype that ameliorated upon a reduction in the endogenous levels of CSPα/DNAJC5 [24]. To test that notion in mice, we examined lipofuscinosis in CSPα-L115R transgenic mice in the homozygous and heterozygous CSPα/DNAJC5 KO background and found that lipofuscinosis persists even under a long-lasting (8 months) 50% reduction of endogenous CSPα/DNAJC5 levels, suggesting that the lipofuscinosis we detected is not a consequence of gain of the normal function of the protein. We cannot rule out that hypermorphic phenotypes might appear in mice upon increasing the genetic dose of the mutant transgenes as it occurs in Drosophila. In addition, in our mice models, the promoter Thy1 drives the expression only in neurons but not in microglia, however, since CSPα/DNAJC5 is also expressed in microglia [41], it is conceivable that CLN4 human patients might suffer from microglial dysfunction and that the major phenotype arises when both, neurons and microglia, are dysfunctional [50]. Indeed, it has been proposed that in CLN11 the neurological alterations arise only when different cell types, including neurons and glial cells simultaneously lack progranulin [42–44]. In that sense, the mice generated in the present study do not develop an overt phenotype perhaps because the mutant protein needs to be expressed in neurons and microglia to unfold the full neurological phenotype. In any case, our findings indicate that the neuronal expression of CLN4 mutants is enough to cause neuronal lipofuscinosis that recapitulates human lipofuscinosis by a cell-autonomous mechanism in neurons. In addition, our transgenic models will be advantageous to dissect the specific contribution of neuronal and non-neuronal cell types, such as microglia, in the complex pathology of this disease.

In conclusion, we have developed novel mouse models that, to our knowledge, are the first mammalian models to study the pathological neuronal lipofuscinosis observed in Kufs disease/CLN4 patients. Since using a variety of mouse models, we reveal that the absence of CSPα/DNAJC5 cannot be stated as a primary cause of lipofuscinosis, we hypothesize that the mutant versions of CSPα/DNAJC5 might be interfering with key biochemical pathways such as proteostasis, likely caused by aberrant protein-protein interactions with, so far, unknown molecular partners. That would be a cell-autonomous mechanism occurring through the mutations-mediated gain of a novel, but pathological, function of CSPα/DNAJC5

The mice presented in this study open remarkable possibilities to further investigate the molecular mechanisms of Kufs disease/CLN4 and to test therapeutic strategies to revert pathological lipofuscinosis.

## METHODS

### Mice

All procedures involving animals were performed in accordance with the European Union Directive 2010/63/EU on the protection of animals used for scientific purposes and approved by the Committee of Animal Use for Research at the University of Seville. The animals were maintained in the university animal facilities (IBiS and Macarena campus) on a 12/12-h light/dark cycle with unrestricted access to food and water.

### Generation of transgenic mice

GFP-CSPα fusion protein was excised from pEGFP-C1-CSPα with SmaI and NheI and cloned under control of Thy1 promoter into pThy1.2/pUC18 composed vector [10] in HincII (made blunt) and NheI. Mutant versions of CSPα/DNAJC5 cDNA (L115R and L116Del) were produced by double PCR and cloned into AgeI and SacII sites ot pThy1.2 vector. Primers used for cloning described in Supplementary Table 4. For transgenic mice generation, expression cassette (8222 or 8219bp) was excised with NotI and PvuI sites, to remove bacterial sequences, purified from agarose gel and resuspended in 7.5mM Tris pH7.4, 0.2mM EDTA. Pronuclear injections were carried out by “Centro de Producción y Experimentación Animal” at the University of Seville. Briefly, super-ovulated FVB females were mated and after vaginal plug, pronuclear fertilized oocytes were extracted. DNA fragments were injected into male pronucleus and transferred into pseudopregnant FVB females. Conditional floxed mouse line Dnajc5flox/flox was previously described [36]. For Cre-recombinase induction, mice were fed a tamoxifen-enriched diet (TAM400/CreER, TD.55125.I, Envigo) for 30 days to induce genetic removal of CSPα/DNAJC5. Mice were fed with tamoxifen at 2 months of age, except the mice that were analyzed 8 months after tamoxifen that were fed with tamoxifen when they were one month old. Genotyping protocols are detailed in Supplementary Table 5.

### Immunohistochemistry and immunofluorescence

For brain fixation and collection, mice were anesthetized with 2% 2,2,2-Tribromoethanol (T48402, Sigma-Aldrich). Mice were perfused with Phosphate Buffered Saline (PBS) 1X (137 mM NaCl, 2.7 mM KCl, 10 mM Na2HPO4, 2 mM KH2PO4, pH 7.4) to eliminate blood from tissues, and then with 4% paraformaldehyde (PFA) in PBS to fix tissues. Collected brains were post-fixed in PFA 4% for 24 hours at 4°C and then cryoprotected in 30% sucrose in PB 1X with 0.01% sodium azide. A Leica CM 1950 cryostat was used for serial sectioning of either sagittal or coronal planes of 40 µm thickness. Slices were stored at −20°C in 50% glycerol in PBS 1X until use. For those antibodies that need antigen-retrieval to recognize their targets, free-floating brain slices were incubated in 10 mM sodium citrate pH 6 in PBS during 30 min at 80°C, and then blocked with 2% non-fat milk and 0.3% Triton X-100 in PBS for 30 min. Then, slices were incubated in blocking solution with 3% FBS and 0.3% Triton X-100 in PBS for 1 hour at room temperature. Incubation with primary antibodies (see Supplementary Table 2) was performed overnight at 4°C in the same blocking solution. Secondary antibodies were incubated in same blocking solution in dark for 2 hours at room temperature (see Supplementary Table 3). Slices were then washed, mounted and coverslipped with FluorSave Mounting Medium (#345789. Millipore). In case of immunohistochemistry, endogenous peroxidase was inactivated for 10min in 8%methanol, 2.5% H2O2 in PBS. Biotinylated secondary antibody was used, and kit ABC used for amplification of signal (Vectastain Elite ABC Kit. #PK-6100. Vector Labs). Peroxidase activity was revealed by 0.02% Diaminobenzidine with 0.01% H2O2 and enhanced with 0.04% NiCl2 in phosphate buffer (10 mM Na2HPO4, 2 mM KH2PO4, pH 7.4). Reaction was stopped by washes in phosphate buffer. Finally, sections were placed in gelatinized glass slides and coverslipped with DPX (Sigma). In case of immunohistochemistry for GVB markers, endogenous peroxidase activity was quenched in 0.3% H2O2 in PBS for 12.5 minutes. Sections were incubated with blocking solution containing 5% normal goat serum (Gibco), 2.5% bovine serum albumin (ThermoFisher Scientific), 0.2% Triton-X100 (ThermoFisher Scientific) in PBS for 1 hour at room temperature. Incubation with primary antibodies (see Supplementary Table 2) was performed overnight at 4°C in blocking solution. Sections were incubated with biotinylated secondary antibody (see Supplementary Table 2) diluted in blocking solution for 90 minutes at room temperature, followed by incubation with Vectastain ABC kit components (Vector Laboratories) (1:800 in blocking solution) for 90 minutes at room temperature. Each of these steps was followed by extensive washing in PBS (three times 10 minutes). Signal was developed using the chromogen DAB (DAB Substrate Kit, Vector Laboratories) and sections were washed in H2O (two times quickly, two times five minutes). Sections were counterstained with hematoxylin (diluted 1:1 in MilliQ water) for 30 seconds, extensively washed with MilliQ water and de-stained in acidic ethanol (HCl 1:100 diluted in 70% EtOH) for a few seconds. Sections were mounted on gelatin-coated microscope slides, dehydrated, and mounted using Quick-D mounting medium (Klinipath BV). Stained sections were imaged using a Leica DM2500 microscope with Leica MC170 HD camera. For microglia immunostaining analysis, free-floating brain slices were first rehydrated and then incubated for 2h in PBS containing 0.3% Triton X-100, 10% normal donkey serum, and 1% BSA, followed by incubation with primary antibodies (see Supplementary Methods) overnight at 4°C. After three washes in PBS, sections were incubated with secondary antibodies for 2h at room temperature.

### Acquisition and analysis of confocal images

Confocal images were acquired in a Nikon A1R+ confocal or Leica Stellaris 8 microscopes using immersion 20x/63x objectives. Maximal Z-projection was done choosing the same number of planes for each staining. Quantification was done with the help of custom macros for Fiji (ImageJ) (available at: https://github.com/SLopezBegines/ImageJ-Macros) to quantify size, density and intensity of lipofuscin granules in CA3 hippocampus region. In brief, by using Analyze Particles tool, a specific ROI was done for every particle above of a threshold applied to a selected channel (usually 561 laser line channel) and with a size above of 10 µm^2^. Distribution of ROIs was saved (white panels); and size, density of accumulations in CA3 area and intensity of signal in each ROI in channel was quantified. R^2^ Pearson’s correlation index for colocalization analysis was obtained using Coloc2 3.0.5 plugin from ImageJ. 3D reconstruction was performed using Bitplane Imaris x64 9.6.0 image analysis software (Oxford Instruments, Concord, MA). Images were first subjected to a background subtraction and then processed with the surface and filament module to rebuild microglia, cell bodies, microglia processes, lipofuscin particles and neurons.

### Transmission electron microscopy

Brains were collected after anesthesia and sectioned at 200 µM thickness using vibratome (Leica VT1200S) immersed in ice cutting solution (222 mM sucrose, 11 mM glucose, 3 mM KCl, 1 mM NaH2PO4, 26 mM NaHCO3, 7 mM MgCl2 and 0.5 mM CaCl2 in ddH2O). Sections were then fixed in a 0.1 M sodium cacodylate pH 7.4, 2.5% glutaraldehyde solution for 2 hours at room temperature, followed by washing with 0,1 M cacodylate twice for 10 minutes. Sections were kept in 0.1 M cacodylate with 0.01% sodium azide at 4°C until use. For Spurr resin inclusion, samples were fixed in 1% OsO4, stained with 2% Uranyl acetate, dehydrated in a gradient up to 100% acetone and infiltrated in a gradient of acetone to Spurr resin. Ultrathin 70nm sections were observed in a Zeiss Libra 120 transmission electron microscope. To quantification, number of lipofuscin accumulations was quantified in each 2kx field and average number of accumulations per image and per mouse was used. Numbers describe number of images and number of mice used.

### Immunoblotting and quantification

Fresh tissue was flash-frozen in liquid nitrogen and kept at −80C until use. Hippocampal samples were homogenized in Homogenization Buffer (50mM Tris-HCl pH 7.4, 150mM SDS, 1%SDS, 5mM EDTA, 1mM glycerophosphate, 1 mM NaF, 1mM PMSF, 1 complete protease inhibitor cocktail tablet #11697498001. Merck) with blue pestles and then under rotation for 2h in cold-room, followed by centrifugation at 16000g for 30min at 4°C. Protein content of soluble fractions were quantified using Pierce BCA Protein Assay Kit (23227, Thermo Scientific), following manufacturer’s instructions and normalized. Samples were boiled in Laemmli 1X (50 mM Tris-HCl, pH 6.8, 10% glycerol, 2% SDS, 0.0067% bromophenol blue, 0.1 M α-mercaptoethanol), and boiled at 95°C for 5 minutes. Samples were resolved in standard 12% Tris-glycine polyacrylamide gels and transferred to PVDF membranes. Protein signal was detected using enhanced chemiluminescence (ClarityTM Western ECL Substrate, 170-5060, Biorad) and images were taken using the Chemidoc Touch Imaging System (BioRad) and analyzed with Fiji (ImageJ). Data were analyzed using Microsoft Excel and GraphPad Prism 8.

### Data availability

Data supporting the findings of this study are available from the corresponding author.

## ACKNOWLEDGEMENTS

This work was supported by the Spanish Agencia Estatal de Investigación and Ministerio de Ciencia e Innovación (BFU2016-76050-P, PID2019-105530GB-I00/AEI/10.13039/501100011033, BES-2011-046029, BES-2017-082324), the Andalusian Department of University, Research and Innovation (CUII, P12-CTS-2232, P18-FR-2144, CTS-600), the Andalusian Department of Health and Consumption (RH-0064-2021), the Institute of Health Carlos III (ISCIII) and European Regional Development Fund (ERDF) to RFC; ZonmMW Memorabel / Alzheimer Nederland #733050101 to WS; and PID2021-125875OB-I00, MCIN/AEI/ 10.13039/501100011033, “ERDF A way of making Europe”, and Junta de Comunidades de Castilla-La Mancha (SBPLY/21/180501/000064) to RL. We are grateful to Dr. Sreeganga Chandra members from her lab for critical reading of a previous version of the manuscript and insightful comments, and to staff members at research facilities of IBiS and Centro de Investigación, Tecnología e Innovación de la Universidad de Sevilla (CITIUS) for technical support and advice. Thanks to A. Arroyo and M.C. Rivero for previous technical assistance with genotyping. We are indebted to Dr. Oscar Pintado for his pioneering work in the generation of genetically modified mice, including mice reported in this study, during more than two decades at the University of Seville.

## AUTHOR CONTRIBUTIONS

S.L.-B generated and characterized mouse lines, designed and performed experiments and analyzed data; A.L.-R and F.M. designed and carried out plasmid constructions for transgenesis; A.L.-R. carried out generation and characterization of transgenic mouse lines; C.M.-C generated a conditional knock-out mouse line and performed experiments and analyzed data; N.B. designed experiments and analyzed data pertaining to microglia; V.W. and W.S. performed experiments and analyzed data pertaining to GVB; S.L.-B, C.A and R.L.-M performed experiments and analyzed data pertaining electron microscopy data; J.L.N.-G set-up protocols for electron microscopy and co-supervised C.M.-C experiments; R.F.-C. conceived the study, designed experiments, analyzed data and wrote the paper with input from all authors.

## CONFLICT OF INTEREST

The authors declare no conflict of interest.

## SUPPLEMENTARY INFORMATION

### SUPPLEMENTARY FIGURES

**Supplementary Figure 1.**
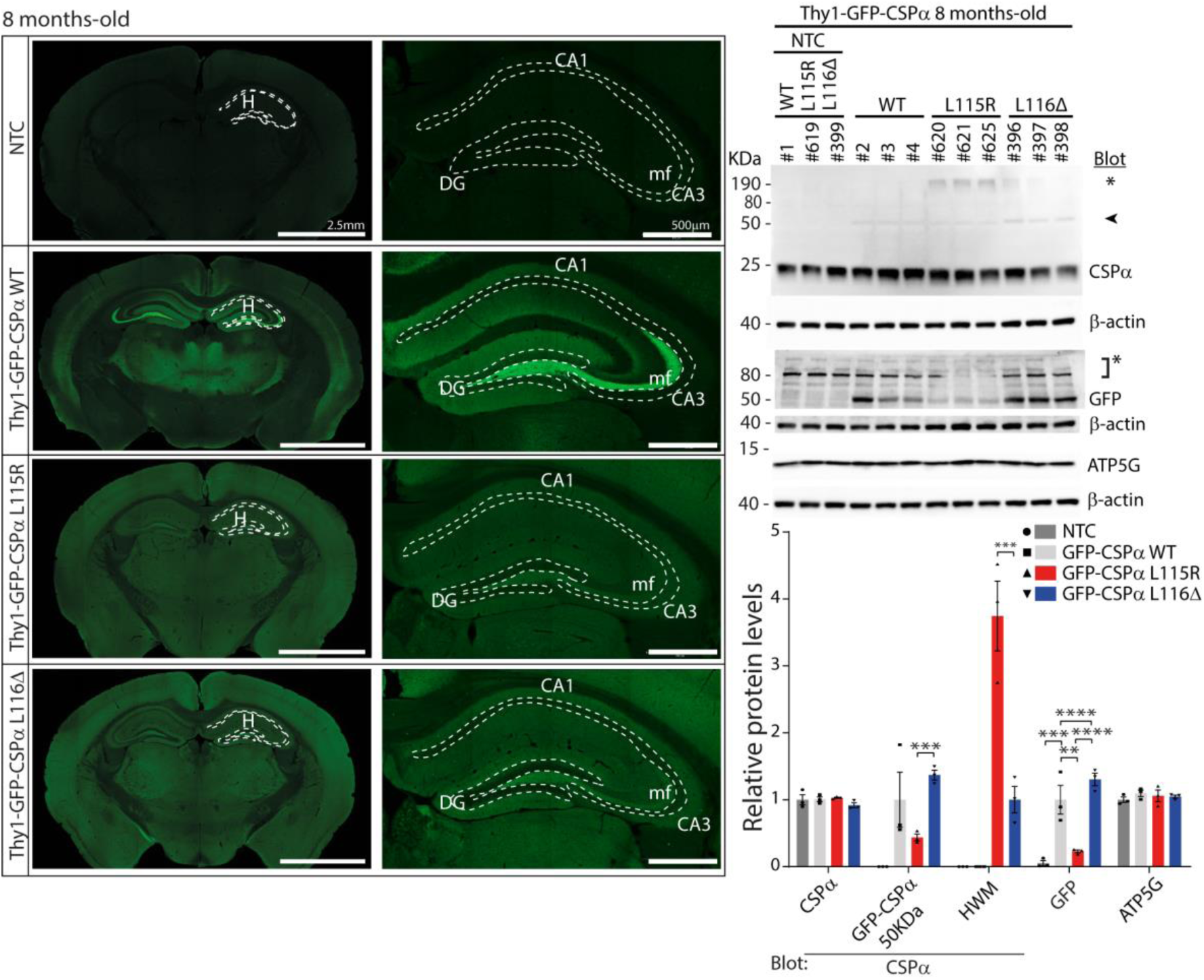
Transgenic expression of wild-type and CLN4 mutant forms of CSPα/DNAJC5 in mouse brain at 8months of age. **A**. Representative epifluorescence images of GFP immuno-staining demonstrate widely distributed expression of all transgenes in brain in 8 months old mice. General stronger transgene expression of GFP-CSPα WT especially evident at hippocampal mossy fibers. No signal is detected in non-transgenic control (NTC) mice. Scale bar 500µm. **B**. Transgenic proteins detected by western blot of hippocampal extracts from 8 months old mice. Numbers indicate mouse ID number. Upper blot, endogenous CSPα/DNAJC5 is detected in all samples while a band corresponding to GFP-tagged CSPα/DNAJC5 (arrow head, 50KDa) appears in transgenic samples but not in NTC. High molecular weight species (asterisk *) are detected in mutant transgenic samples, especially in the L115R mutant. GFP signal is only detected in transgenic samples. Non-specific band due to GFP antibody (asterisk *). **C**. Levels of selected hippocampal proteins. Relative protein levels normalized to non-transgenic mouse lines, except for GFP quantification that was normalized to GFP levels of the GFP-CSPα-WT transgenic line. Quantitative data available at Supplementary Table 1. Two-way ANOVA with Tukey’s post hoc test (*p<0.05, \*\**p*<0.01, \*\*\**p*<0.001*, ****p<0.0001)*.

**Supplementary Figure 2.**
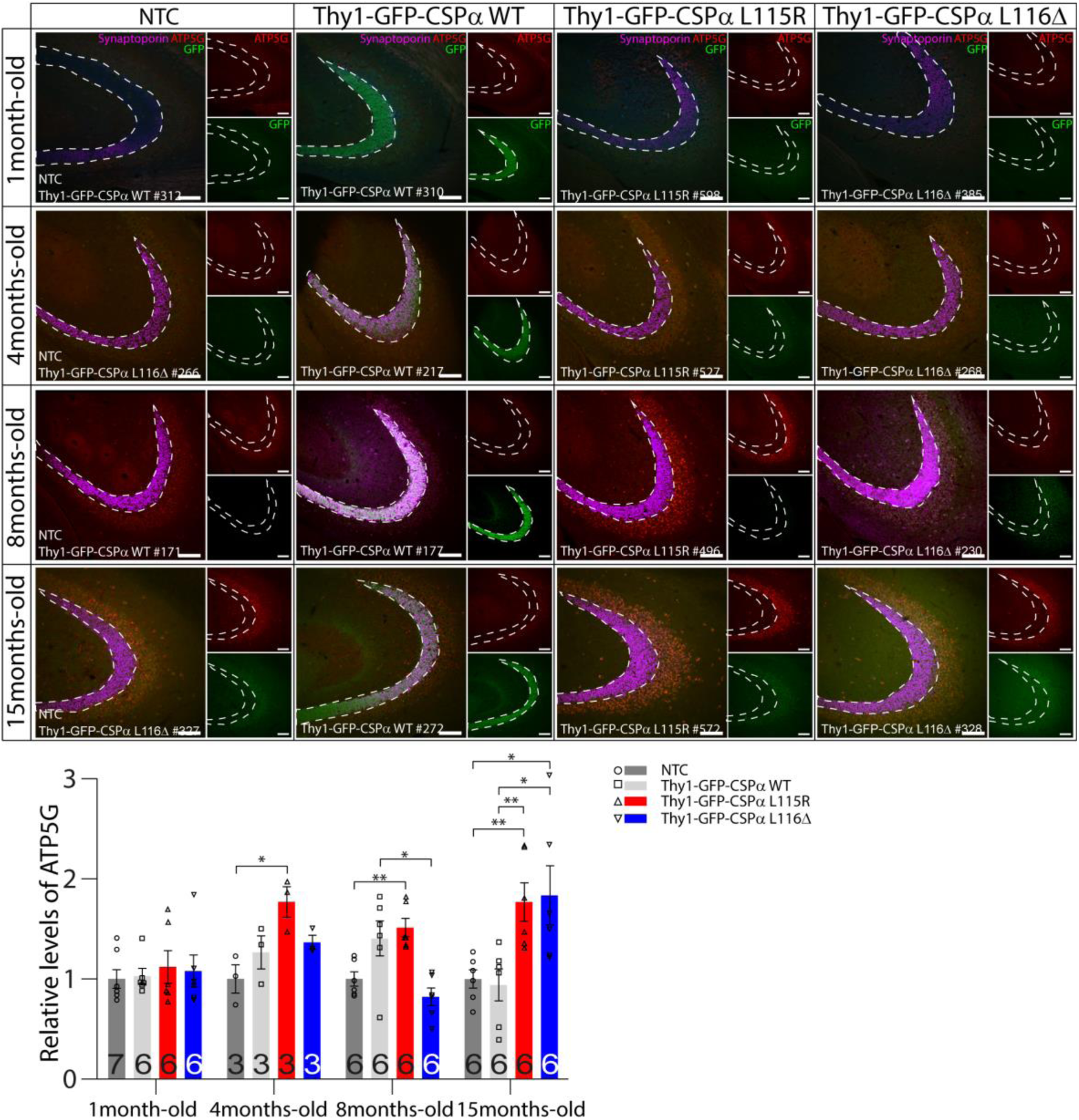
Preferential detection of ATP5G signal at the CA3 region after 4 months of age in CLN4 mutant mice. Immunolabeling of pyramidal neurons at the CA3 hippocampal region with antibodies against ATP5G (red) and GFP (green) and presynaptic mossy fiber with an antibody against synaptoporin (magenta) from 4 different genotypes (non-trasngenic control, GFP-CSPα-WT, GFP-CSPα-L115R and GFP-CSPα-L116Δ) at different ages (1, 4, 8 and 15 months-old). Mice identification numbers shown in every image. Number at bottom of graph bars indicates number of images from 3 mice per each genotype. Scale bar 100µm. Unpaired t-test *p<0.05, **p<0.01. Quantitative data available at Supplementary Table 1.

**Supplementary Figure 3.**
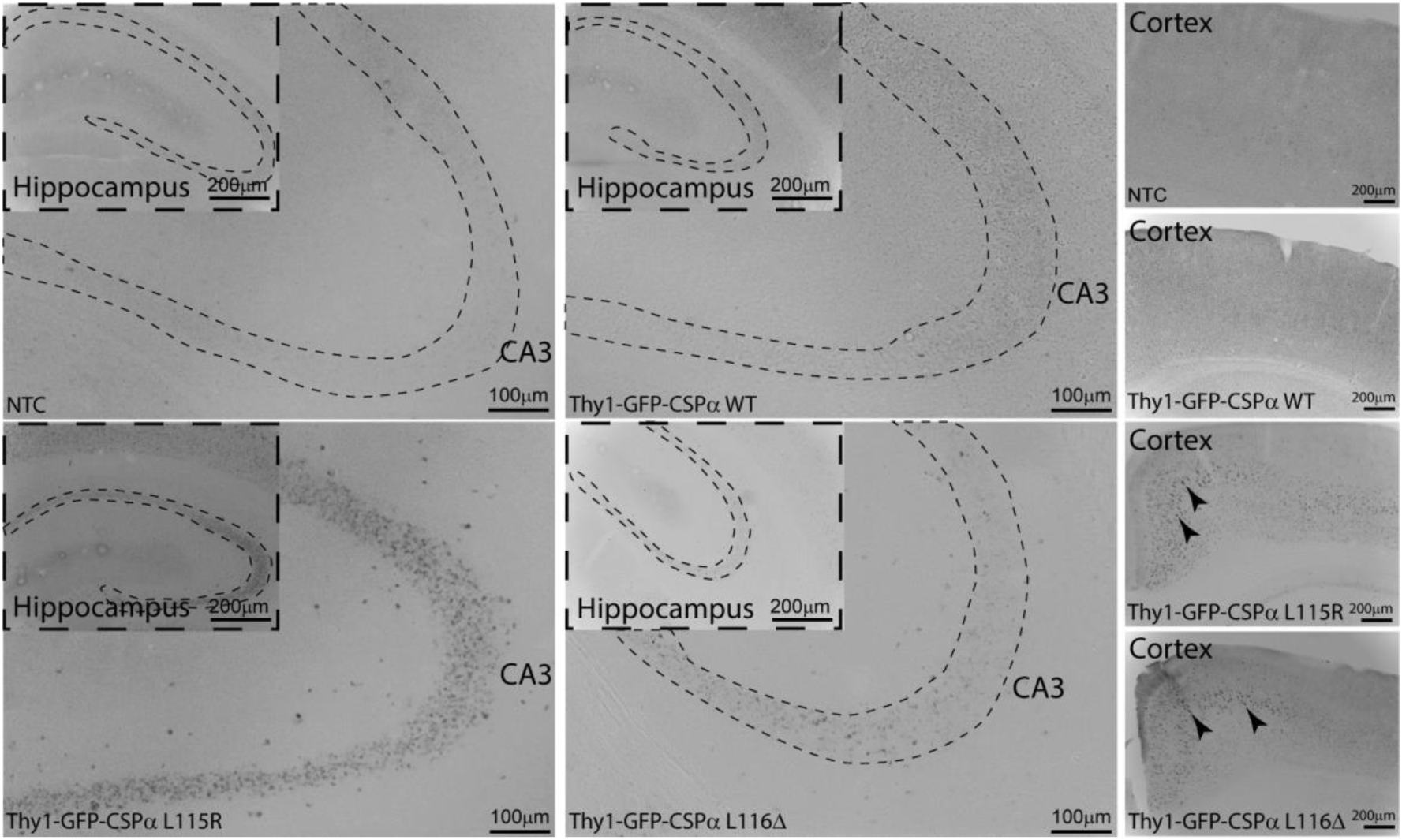
Pathological lipofuscinosis visualized with bright field microscopy. Immunohistochemical labeling based on peroxidase-staining demonstrate the pathological presence of ATP5G accumulations at the CA3 region of hippocampus and cortex in Thy1-GFP-CSPα-L115R and Thy1-GFP-CSPα-L116Δ mice but not in controls.

**Supplementary Figure 4.**
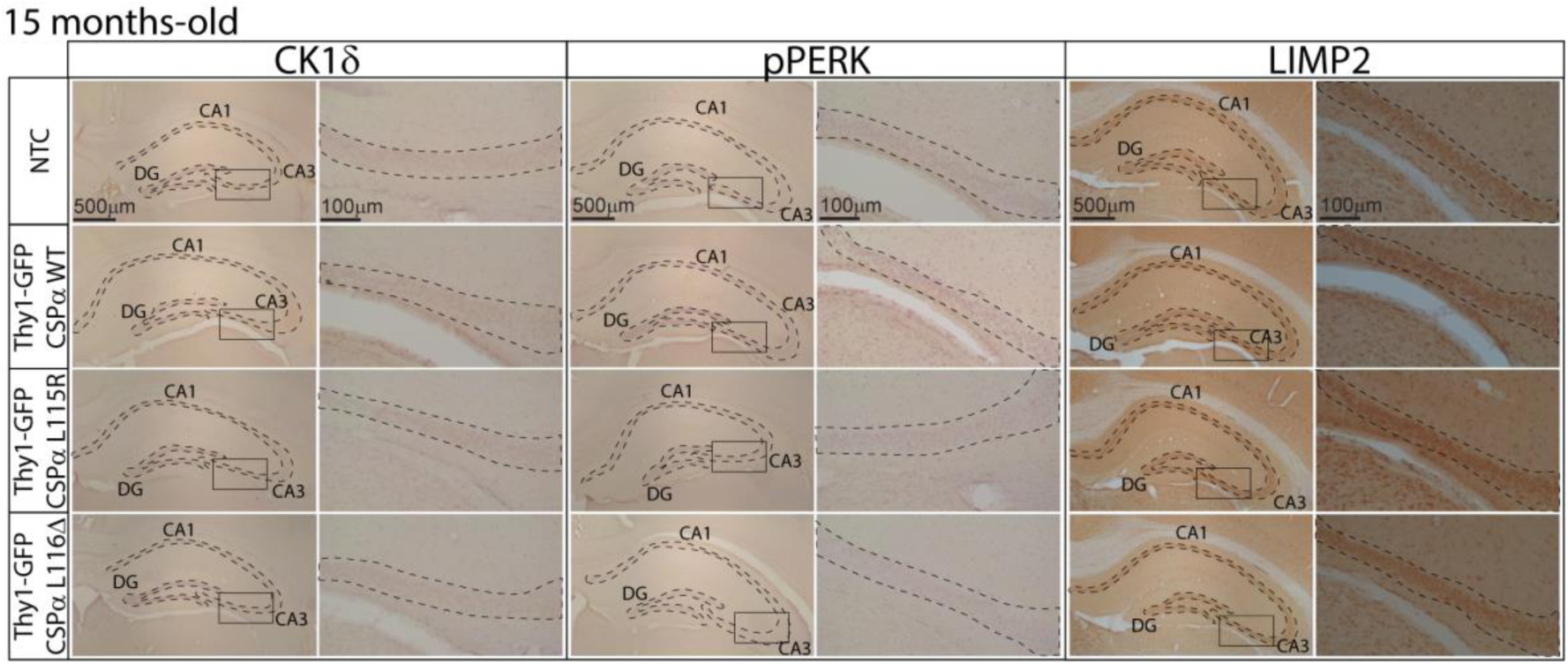
CLN4 CSPα/DNAJC5 mutations do not lead to hippocampal granulovacuolar degeneration body (GVB) formation. Immunohistochemical labeling with antibodies against CK1δ, pPERK and LIMP2 of hippocampal sections of non-transgenic control (NTC), Thy1-GFP-CSPα-WT, Thy1-GFP-CSPα-L115R and Thy1-GFP-CSPα-L116Δ in 15 months-old mice. No GVBs are found in mice from any of the genotypes, as shown by immunolabeling of CK1δ and pPERK using antibodies that detect GVBs in mice with tau pathology. There is no obvious difference in immunolabeling with the lysosomal and GVB membrane marker LIMP2 detected between genotypes. Nuclei are counterstained with hematoxylin in all images.

**Supplementary Figure 5.**
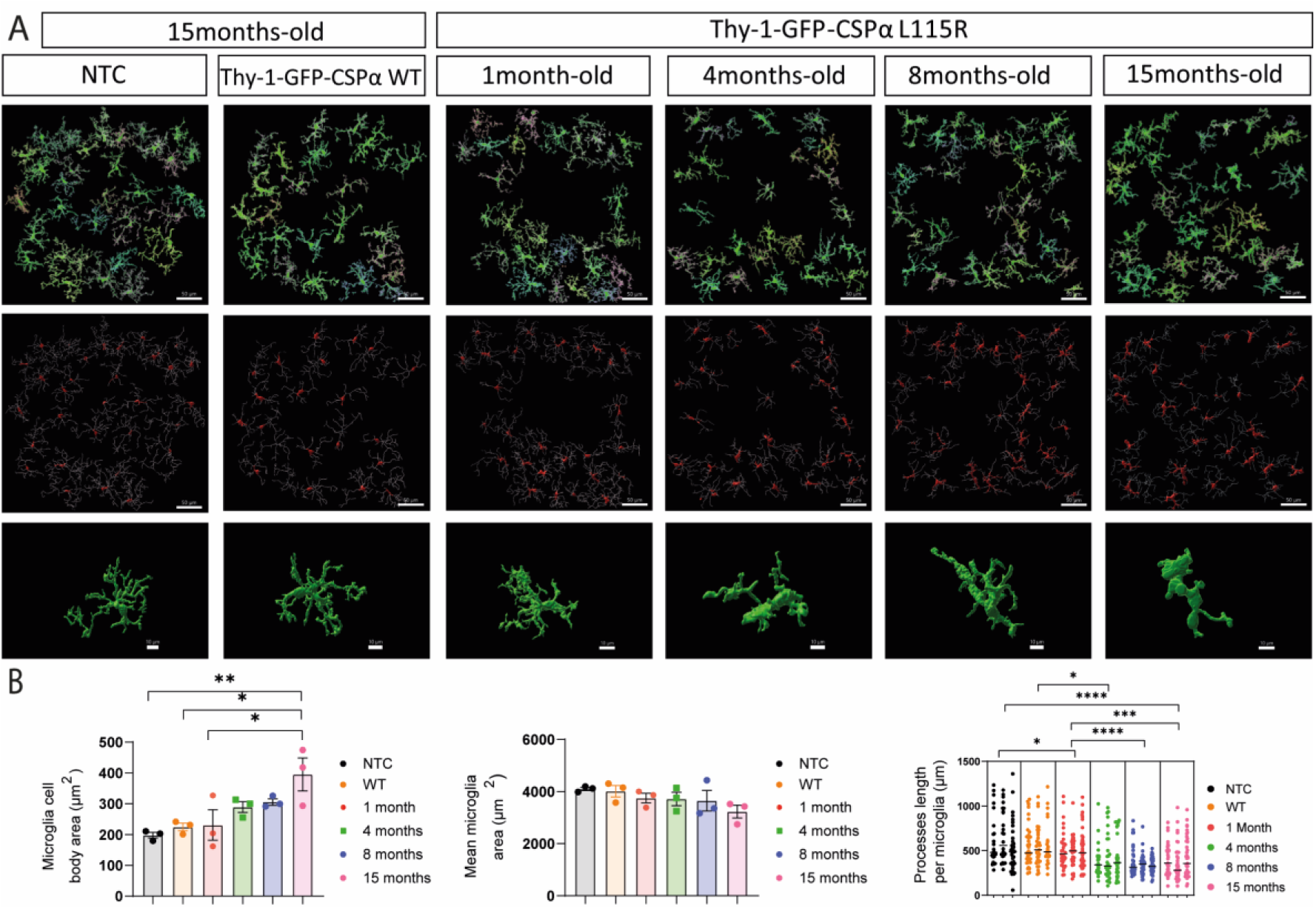
Morphological analysis of microglia in Thy1-GFP CSPα L115R transgenic mice. **A. First row**, representative images of immunostaining of the CA3 region microglia (Iba-1, green) in control (NTC) and GFP-CSPα-WT (both at 15 months of age) and 1-, 4-, 8-and 15 months old GFP-CSPα-L115R mice; Scale bar 50µm. **Second row**, Imaris 3D Reconstruction for the analyzed cells; Scale bar 50µm. **Third row**, Imaris cell bodies and filament reconstruction of the analyzed cells; Scale bar 50µm. **Bottom row**, Imaris 3D reconstruction of exemplary individual microglia cells; Scale bar 10µm. **B**. Quantification of cell body, mean area and mean processes length of microglia in NTC and GFP-CSPα-WT (both at 15 months of age), and GFP-CSPα-L115R at 1-, 4-, 8- and 15-months old mice were represented. Data were presented as mean ± SEM, two-way ANOVA with Tukey’s post hoc test (*p<0.05, **p<0.01, ****p<0.0001), N=3 mice/group, 3 images/mouse. At least 25 microglia cells/per mouse were analyzed to get the processes length of microglia data. Quantitative data available at Supplementary Table 1. NTC, Thy1-GFP-CSPα-WT and Thy1-GFP-CSPα-L115R representative images are the same displayed in Fig. 4. Different images from the same NTC and Thy1-GFP-CSPα WT set of mice were used in Supp. Fig. 5 and Supp. Fig. 6 for morphological analysis of microglia.

**Supplementary Figure 6.**
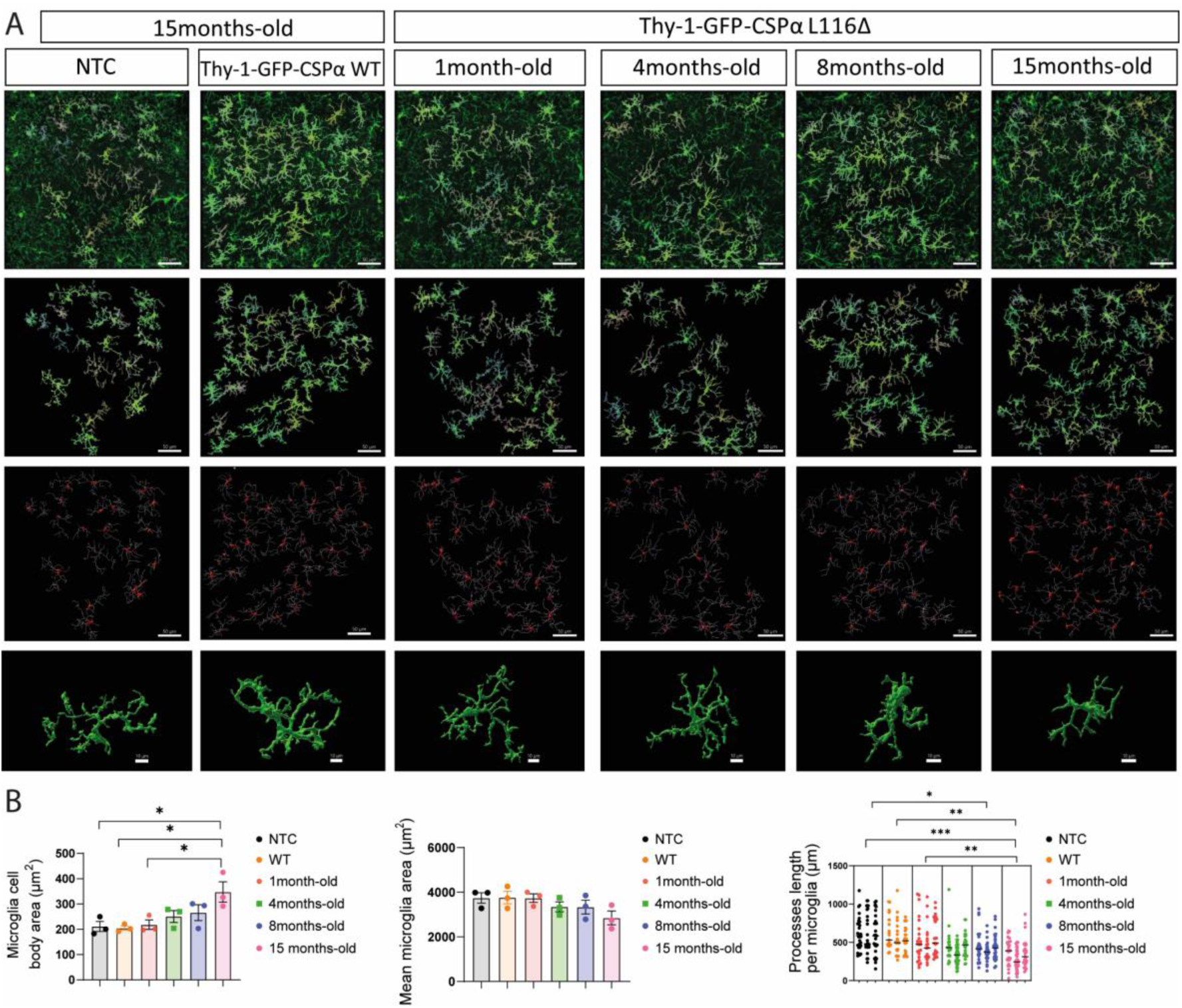
Morphological analysis of microglia in Thy1-GFP CSPα L116Δ transgenic mice. **A. First row**, representative confocal images of immunostaining of the CA3 region microglia (Iba-1, green) of old control (NTC) and GFP- CSPα-WT (both at 15 months of age), and GFP-CSPα- L116Δ mice (at 1, 4, 8 and 15 months of age). Scale bar 50µm. **Second row,** Imaris 3D reconstruction for the analyzed cells. **Third row**, cell bodies and filament reconstruction of the different cells analyzed using Imaris software. Scale bar 50µm.**Bottom row**, Imaris 3D reconstruction of exemplary individual microglia cells; Scale bar 10µm. **B.**Quantification of cell body, mean area and mean processes length of microglia in NTC and GFP-CSPα-WT (both at 15 months of age), and GFP-CSPα- L116Δ at 1-, 4-, 8- and 15-months old mice, were represented. Scale bar 50µm.. Data were presented as mean ± SEM, Two-way ANOVA with Tukey’s post hoc test (*p<0.05, **p<0.01), N=3 mice/group, 3 images/mouse. At least 20 microglia cells/per mouse were analyzed to get the processes length of microglia data. Quantitative data available at Supplementary Table 1. Different images from the same NTC and Thy1-GFP-CSPα WT set of mice were used in Supp. Fig. 5 and Supp. Fig. 6 for morphological analysis of microglia.

**Supplementary Figure 7.**
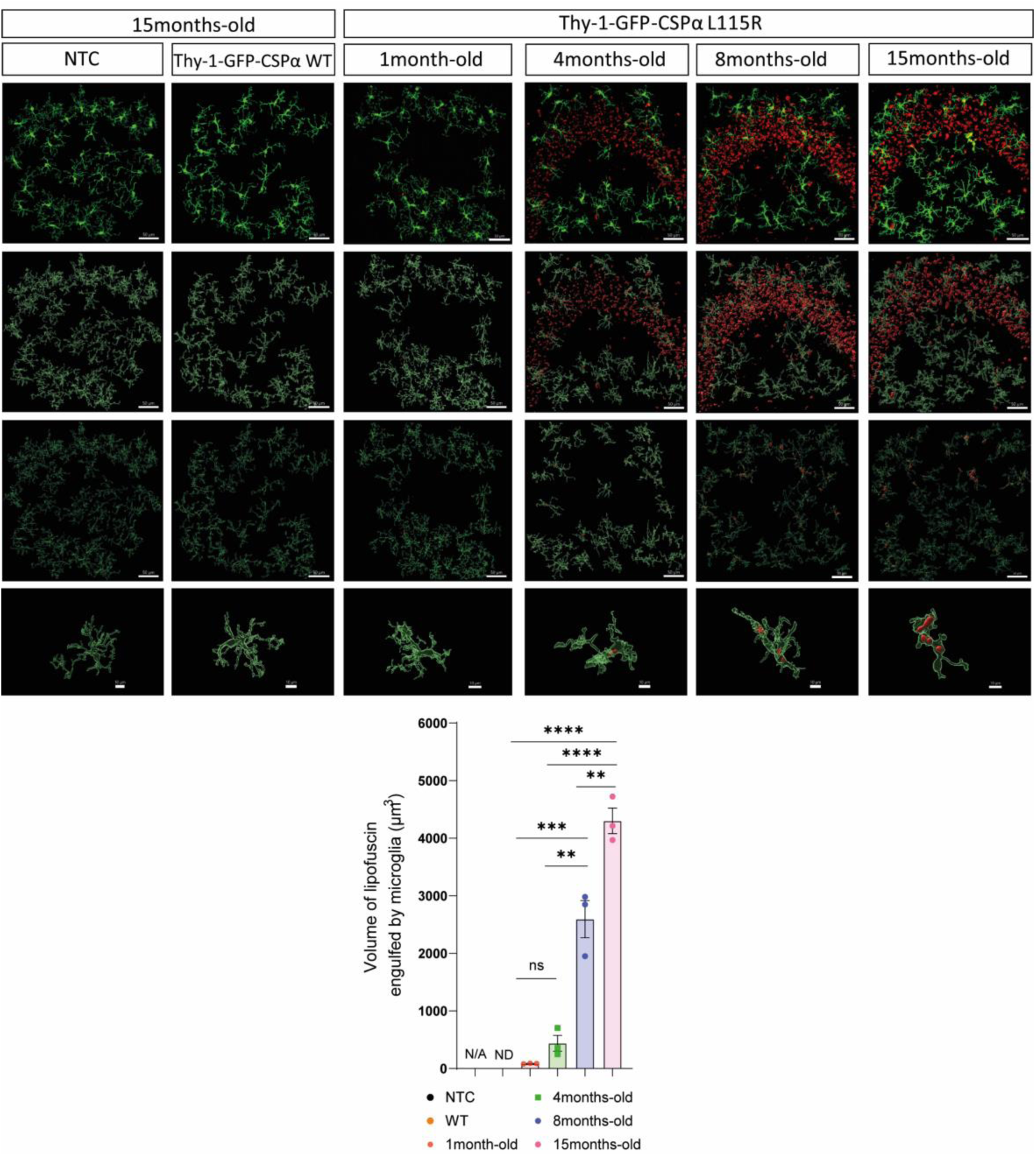
Microglia engulf lipofuscin in Thy1-GFP CSPα L115R transgenic mice. Upper panel. **First row**, representative confocal immunostaining images of microglia (Iba-1, green) and lipofuscin (red, autofluorescence, TRITC filter) at the CA3 region of 1, 4, 8 and 15 months old Thy1-GFP CSPα L115R mutant mice. **Second row**, Imaris 3D reconstruction. **Third row,** Imaris 3D reconstruction of lipofuscin engulfed by microglia. Scale bar 50µm.**Bottom row,** Imaris 3D reconstruction of exemplary individual microglia cells engulfing lipofuscin; Scale bar 10µm. **Lower panel.** Quantification of volume of lipofuscin engulfed by microglia in control (NTC) and GFP- CSPα-WT (both at 15 months of age), and GFP-CSPα-L115R at different time points (1, 4, 8 and 15 months old). Data were presented as mean ± SEM, Two-way ANOVA with Tukey’s post hoc test (**p<0.01, ***p<0.001, ****p<0.0001), N=3 mice/group. Different images from the same NTC and Thy1-GFP-CSPα WT set of mice were used in Supp. Fig. 7 and Supp. Fig. 8 for analysis of microglia engulfing of lipofuscinosis. Quantitative data available at Supplementary Table 1.

**Supplementary Figure 8.**
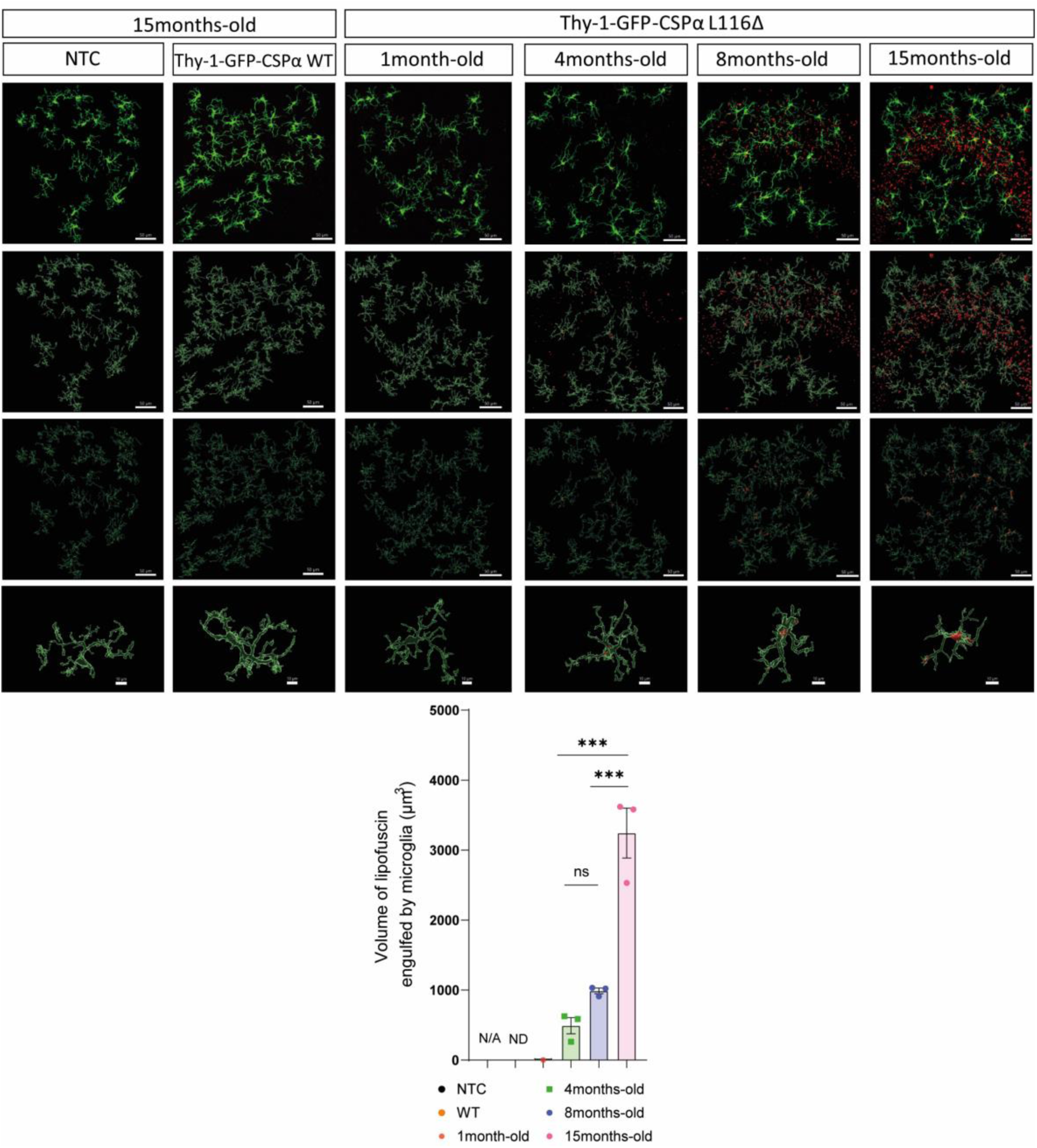
Microglia engulf lipofuscin in Thy1-GFP CSPα L116Δ transgenic mice. Upper panel. **First row**, **r**epresentative immunostaining images of microglia (Iba-1, green) and lipofuscin (red, autofluorescence at 561nm using the TRITC filter) at the CA3 region of 1, 4, 8 and 15 months old Thy1-GFP CSPα L116Δ mutant mice. **Second row**, Imaris 3D reconstruction of lipofuscin and microglia cells. **Third row**, Imaris 3D reconstruction of lipofuscin engulfed by microglia. Scale bar 50µm.**Bottom row,** Imaris 3D reconstruction of exemplary individual microglia cells engulfing lipofuscin; Scale bar 10µm. **Lower panel.** Quantification of volume of lipofuscin engulfed by microglia in 15 months old control (NTC) and GFP-CSPα-WT, and 1, 4, 8 and 15 months old GFP-CSPα-L116Δ mice. Data were presented as mean ± SEM, Two-way ANOVA with Tukey’s post hoc test (***P<0.001,), N=3 mice/group. Different images from the same NTC and Thy1-GFP-CSPα WT set of mice were used in Supp. Fig. 7 and Supp. Fig. 8 for analysis of microglia engulfing of lipofuscinosis. Quantitative data available at Supplementary Table 1.

**Supplementary Figure 9.**
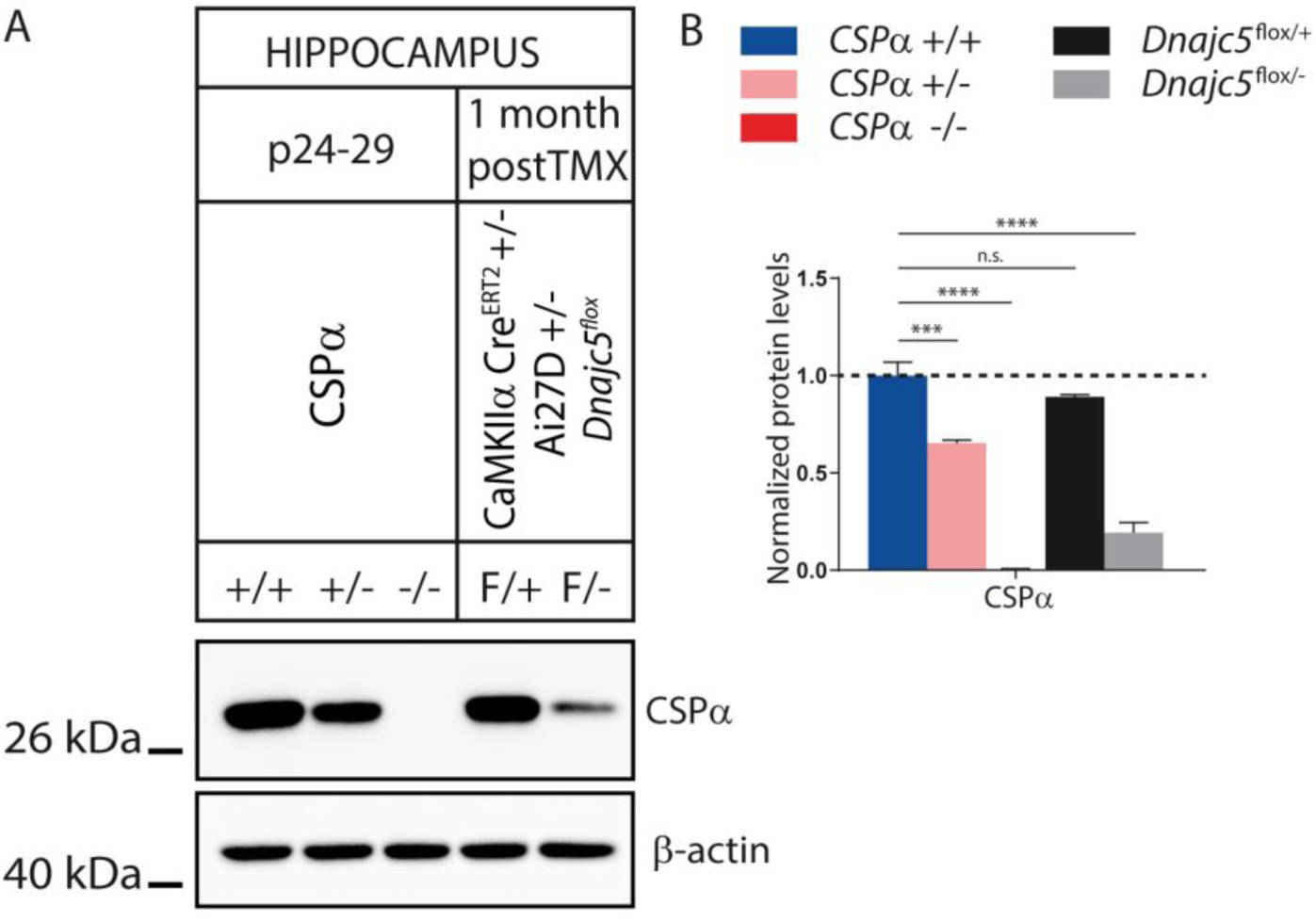
Reduced CSPα/DNAJC5 protein levels in hippocampus in conventional and in glutamatergic-specific conditional CSPα/DNAJC5 KO and heterozygous mice. A. Western blot of hippocampal extracts obtained from CSPα/DNAJC5 WT (+/+), heterozygous KO (+/-) and homozygous KO (-/-) mice at P24- P29 postnatal age and from 3-months-old CaMKIIα^CreERT2^:Ai27D:Dnajc5^flox/+^ and CaMKIIα^CreERT2^:Ai27D:Dnajc5^flox/-^ mice at 1 month after completing the tamoxifen-treatment. B. Quantitation of protein levels using beta-actin as a loading control reveals, as expected, reduction in the levels of CSPα/DNAJC5 in CSPα/DNAJC5 KO and heterozygous mice and in conditional CaMKIIα^CreERT2^:Ai27D:Dnajc5^flox/-^ mice. Quantitative data available at Supplementary Table 1.

**Supplementary Figure 10.**
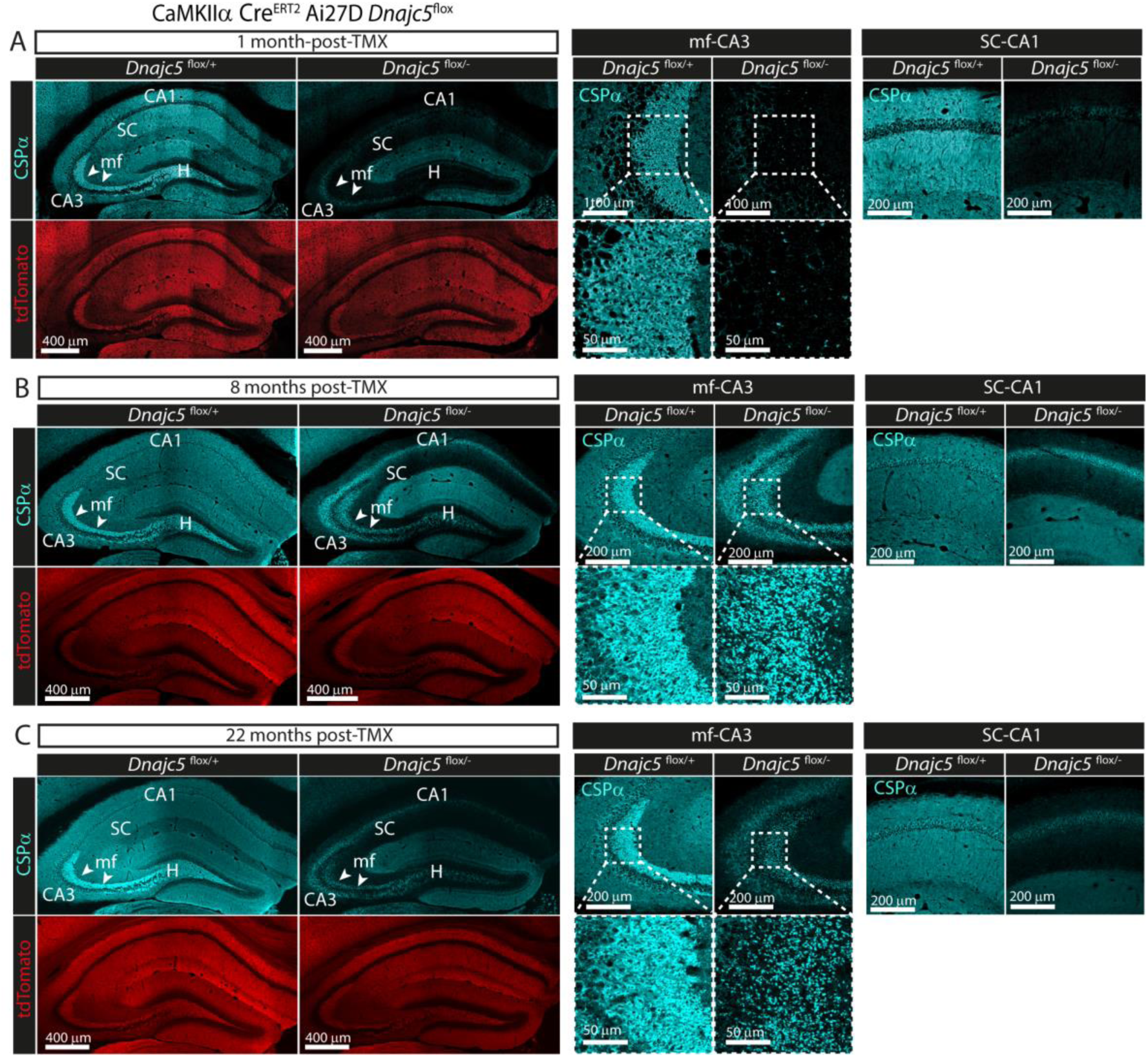
Selective genetic removal of CSPα/DNAJC5 from hippocampal glutamatergic neurons upon tamoxifen administration to CaMKIIα^CreERT2^:Ai27D:Dnajc5^flox/-^ mice A. Three-months old CaMKIIα^CreERT2^:Ai27D:Dnajc5^flox/+^ and CaMKIIα^CreERT2^:Ai27D:Dnajc5^flox/-^ mice were fed with tamoxifen during 30 days and analyzed afterwards at 1-month post-tamoxifen (TMX) by immunolabeling with fluorescently labeled antibodies. Hippocampal sections from CaMKIIα^CreERT2^:Ai27D:Dnajc5^flox/+^ and CaMKIIα^CreERT2^:Ai27D:Dnajc5^flox/-^ conditional KO mice stained with antibodies against CSPα/DNAJC5 (cyan) and against tdTomato (red). Left panel, remarkable decrease of CSPα/DNAJC5 expression in pyramidal neurons at hippocampal CA1, CA3 regions and mossy fibers (mf). Right panel, magnifications of mf- CA3 and SC (Schaffer collaterals)-CA1 synaptic layers reveals the strong decrease in CSPα/DNAJC5 levels. B. As in A, but mice were analyzed at 8-months post-tamoxifen (TMX). In contrast to the results obtained in mice analyzed 1-month post-tamoxifen (TMX), the CSPα/DNAJC5 punctate staining at the stratum lucidum in the CA3 region likely corresponds to novel mossy fibers originated from granule cells that were born through adult neurogenesis once the tamoxifen diet was discontinued. Cre-recombinase was never activated in these new granule cells and therefore CSPα/DNAJC5 expression proceeded normally. C. As in A, but mice were fed with tamoxifen at two months of age and analyzed at 22-months post-tamoxifen (TMX). As in B, CSPα/DNAJC5 expression is detected in a subpopulation of mossy fibers coming from newly born granule cells.

### SUPPLEMENTARY METHODS

**Supplementary Table 2.** Primary antibodies used for immunofluorescence and for western blot.

| *Antigen* | *Reference* | *Concentration used for immunofluorescence* | *Concentration used for western blot* |
| --- | --- | --- | --- |
| CSPα/DNAJC5 | ADI-VAP-SV003-E. Enzo Life Sciences | 1:500 | 1:10000 |
| GFP | #GFP-1020. Aves Lab | 1:500 | 1:1000 |
| Synaptoporin | #102011 Synaptic System | 1:500 | X |
| ATP5G | #ab96655 Abcam | 1:1000<br>Antigen Retrieval | 1:5000 |
| SNAP25 | #836301 SMI81 BioLegend. | 1:5000 | 1:10000 |
| IBA1 | Ab178847 Abcam | 1:500 | N/A |
| Neun | MAB377 Millipore | 1:500 |  |
| Neun | #ABN78 Millipore | 1:500 | N/A |
| β-actin | #A2228 Sigma Aldrich | N/A | 1:10000 |
| Hsc70 | #149011 Synaptic System | 1:500 | 1:2000 |
| tdTomato | #CPCA-mCherry EnCor Biotech | 1:1000 | N/A |
| pPERK | #sc-32577 Santa Cruz | 1:800 | N/A |
| CK1δ | #PA5-32129 ThermoFisher | 1:400 | N/A |
| LIMP2 | #NB400-129 Novus Biologicals | 1:500 | N/A |

**Supplementary Table 3.** Secondary antibodies used for immunofluorescence and for western blot.

| **Secondary antibody** | **Reference** | ***Concentration used for immunofluorescence*** | ***Concentration used for western blot*** |
| --- | --- | --- | --- |
| Anti-Rabbit Alexa Fluor 488 | #711-545-152<br>Jackson ImmunoResearch | 1:500 | N/A |
| Anti-Rabbit Cy3 | #711-165-152<br>Jackson ImmunoResearch | 1:500 | N/A |
| Anti-Rabbit Alexa Fluor 647 | #111-605-144<br>Jackson ImmunoResearch | 1:500 | N/A |
| Anti-Rabbit Biotin-SP | #.711-065-152<br>Jackson ImmunoResearch | 1:1000 | N/A |
| Anti-Mouse Alexa Fluor488 | #715-545-151<br>Jackson ImmunoResearch | 1:500 | N/A |
| Anti-Mouse Cy3 | #715-165-151<br>Jackson ImmunoResearch | 1:500 | N/A |
| Anti-Mouse Alexa Fluor 647 | #115-605-146<br>Jackson ImmunoResearch | 1:500 | N/A |
| Anti-Mouse Alexa Fluor 790 | #115-655-146<br>Jackson ImmunoResearch | 1:500 | N/A |
| Anti-Chicken Alexa Fluor488 | #703-545-155<br>Jackson ImmunoResearch | 1:500 | N/A |
| Rhodamine Red anti Chicken | #703-295-155<br>Jackson ImmunoResearch | 1:500 | N/A |
| Alexa Fluor 647 anti-Chicken | #703-605-155<br>Jackson ImmunoResearch | 1:500 | N/A |
| Anti-Mouse HRP | #115-035-166<br>Jackson ImmunoResearch | N/A | 1:10000 |
| Anti-Rabbit HRP | #111-035-144<br>Jackson ImmunoResearch | N/A | 1:10000 |
| Anti-Chicken HRP | #103-035-155<br>Jackson ImmunoResearch | N/A | 1:10000 |

**Supplementary Table 4.**
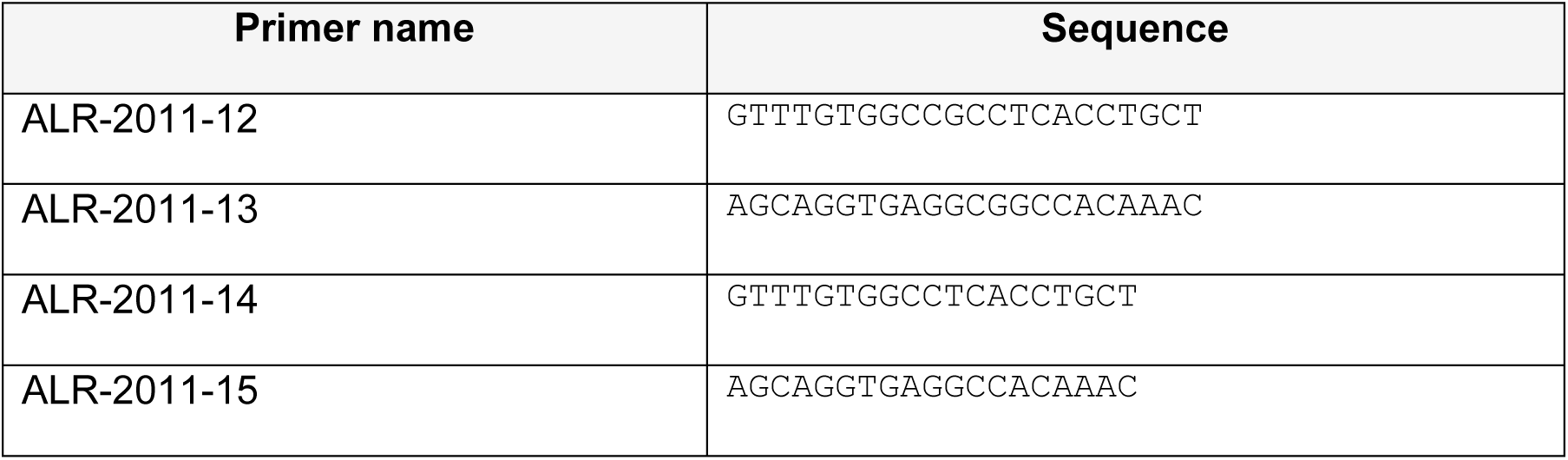
Primers used for cloning.

**Supplementary Table 5.**
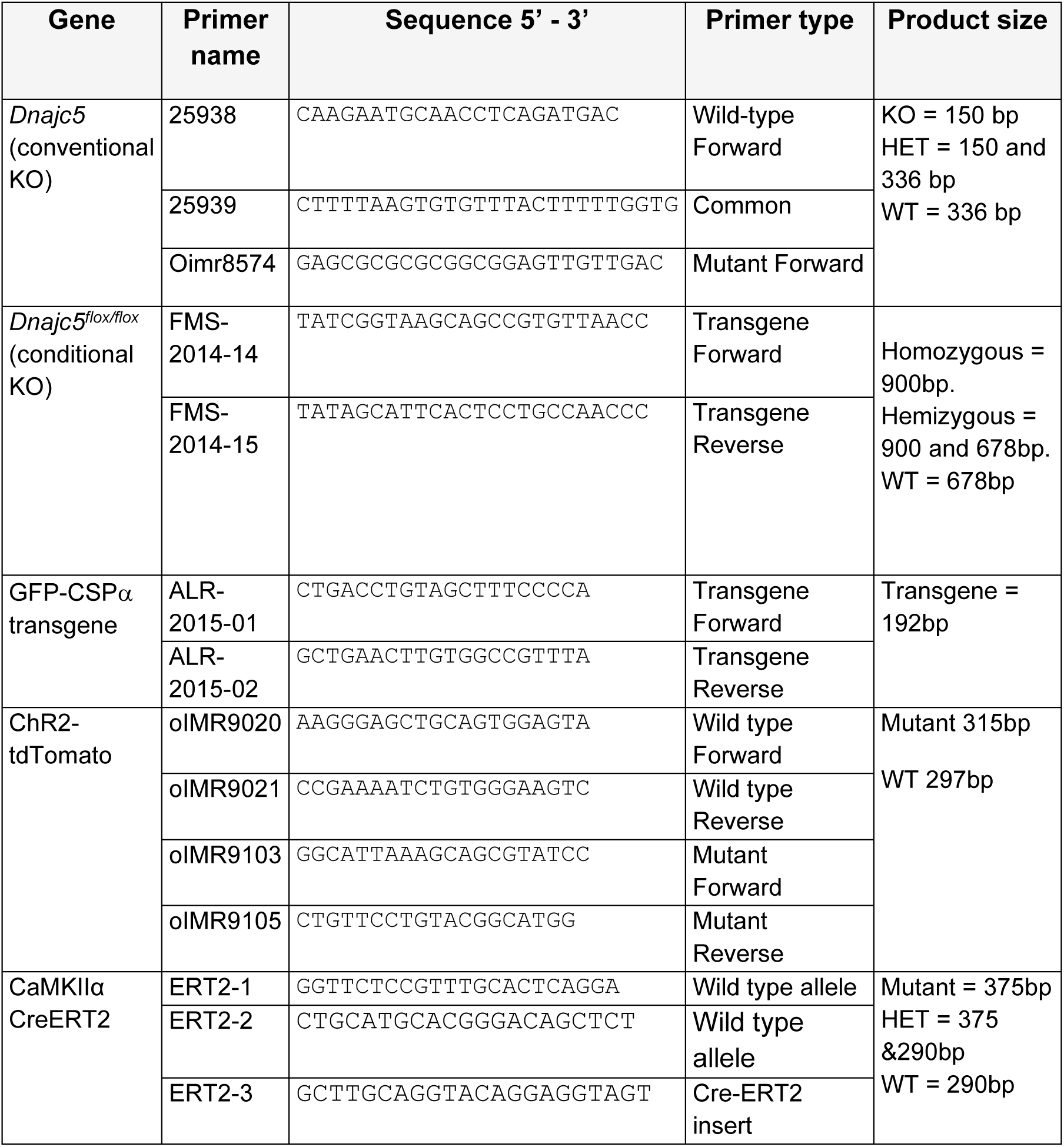
Primers used for genotyping.

## Notes

### Competing Interest Statement

The authors have declared no competing interest.

